# Significant control of Zika infection in macaques depends on the elapsing time after dengue exposure

**DOI:** 10.1101/625293

**Authors:** Crisanta Serrano-Collazo, Erick X. Pérez-Guzmán, Petraleigh Pantoja, Mariah A. Hassert, Idia V. Rodríguez, Luis Giavedoni, Vida Hodara, Laura Parodi, Lorna Cruz, Teresa Arana, Melween I. Martínez, Laura White, James D Brien, Aravinda de Silva, Amelia K. Pinto, Carlos A. Sariol

**Affiliations:** Department of Microbiology and Medical Zoology, University of Puerto Rico-Medical Sciences Campus, San Juan, Puerto Rico, United States of America; Unit of Comparative Medicine, Caribbean Primate Research Center, University of Puerto Rico-Medical Sciences Campus, San Juan, Puerto Rico, United States of America; Department of Molecular Microbiology & Immunology, Saint Louis University School of Medicine, Saint Louis, Missouri 63110 USA; Departments of Virology and Immunology, Texas Biomedical Research Institute, San Antonio, Texas, United States of America; University of North Carolina Chapel Hill, NC; Department of Internal Medicine, University of Puerto Rico-Medical Sciences Campus, San Juan, Puerto Rico, United States of America

**Keywords:** Dengue, Zika, T cells, macaques

## Abstract

Prior exposure to a single serotype of dengue virus (DENV) predisposes individuals to severe disease upon secondary heterologous DENV infection. Here we show that the length of time between DENV/Zika (ZIKV) infections has a qualitative impact on controlling ZIKV replication. We identified limited but significant differences in the magnitude of the early humoral immune response associated with a period of twelve months but not three months of DENV convalescence. However, their role limiting ZIKV replication is not conclusive. There was no evidence of in vivo antibody-dependent amplification of ZIKV by DENV immunity in any group. We are also showing that the significant differences among groups may be linked to a pre-existing polyfunctional CD4+ T cells response (increased IFN-g and Cd107a before ZIKV infection) and to an early and continuous expansion of the CD4+ effector memory cells early on after ZIKV infection. Those significant differences were associated with a period of 12 months after DENV infection that were not observed in a span of 3-months. These results suggest that there is a window of optimal cross-protection between ZIKV and DENV with significant consequences. These results have pivotal implications while interpreting ZIKV pathogenesis in flavivirus-experimented populations, diagnostic results interpretation and vaccine designs among others.

**Author Summary:** Since its introduction in the Americas region ZIKV virus has been associated to severe birth defects. One of the questions that remains open is the role of previous dengue or any other flavivirus immunity in the pathogenesis of ZIKV and more important, if the time elapse between DENV and ZIKV play a role enhancing ZIKV pathogenesis as it is the case for subsequent DENV infections. On this work, using NHP as a model we compared the effect of a period of 12 months vs. a period of 3 months of DENV immunity in the outcome of ZIKV infection. We found that previous DENV infection, at any of the tested period of time do not induce ZIKV enhancement. More relevant are showing that when the two infection occurs at least one year apart the preexisting DENV immunity is better at controlling ZIKV replication and that the role of the neutralizing antibodies is very limited. On the contrary our results suggest that early after ZIKV infection the cellular immune response, may plays a predominant role. Our findings have critical relevance to understand the dynamic interaction between these two flavivirus, their pathogenies, diagnosis and vaccine design.

---

Zika virus (ZIKV) spread in the Americas has been linked to unique severe adverse outcomes such as fetal loss (1), congenital Zika syndrome (CZS) (2), Guillain-Barré syndrome (GBS) (3), and rare cases of encephalopathy (4), meningoencephalitis (5), myelitis (6), uveitis (7), and severe thrombocytopenia (8). Previous studies have shown that prior exposure to a single serotype of dengue virus (DENV) predisposes individuals to severe disease upon secondary heterologous DENV infection. However, the association of previous flavivirus exposure at any time before ZIKV infection with the severe adverse outcomes of the infection is still unclear.

ZIKV infection remains a public health concern with the recent epidemic infecting millions of people in the Americas^8^. ZIKV also poses a pandemic threat, with studies demonstrating that a larger range of the tropical and subtropical regions have suitable conditions for ZIKV transmission and dissemination^1^. ZIKV is mainly transmitted through the bite of *Aedes aegypti*, the same vector implicated in DENV infection, there are however other routes of infection including sexual contact (9-11) and vertical transmission (12-14) that increase the spread of the virus.

Diseases associated with arboviruses are cyclical. While viruses like DENV and yellow fever (YFV) remain endemic in multiple areas of the world, outbreaks of increased disease severity associated with these viruses occur intermittently over the course of multiple years. The timing of these outbreaks is particularly relevant for DENV as the time between a primary and a secondary DENV infection is relevant to the clinical presentation. A short interval between homologous or heterologous DENV infection usually results in protection from disease, while an extended period of time is associated with the potential for severe dengue (15-17), due to either cross-reactive antibodies (18, 19) and/or T cells (20-22).

Little is known about the contribution of virus-specific and cross-reacting antibodies or the cellular immune response generated by a primary DENV infection on the viremia and pathogenesis of a secondary ZIKV infection *in vivo* (23). To address the role of prior flavivirus exposure on ZIKV-associated to disease severity we recently showed that a previous DENV infection (>2 years) does not result in an increase in ZIKV viremia or pathogenesis (24). Interestingly, DENV-immune animals showed a non-statistically significant shorter viremic period compared to DENV-naive macaques.

In this work, we examine the contribution of time between primary and heterologous flavivirus exposure to determine if that factor contributes to cross-protection between ZIKV and DENV. We found that the length of time between the DENV and ZIKV infections has a qualitative impact on controlling the ZIKV infection. In this study, as in our prior work we did not observe evidence of ZIKV disease enhancement associated with prior DENV exposure. We also confirmed that the significant differences among groups are mediated by the pre-existence of a robust effector memory T cell (TEM) and cytotoxic activity mainly mediated by CD4^+^ T cells more than qualitative differences in the humoral immune response. However, by conducting a detailed study of the ZIKV-neutralizing titers vs. ZIKV RNAemia at early time points after infection, we were able to determine a possible contribution of the neutralizing antibodies limiting the ZIKV replication at 7 days after infection in the animals with 12 months but not 3 months of DENV immunity or in the control group. Overall, we demonstrated that exposure to ZIKV 12 months after DENV infection afford a high level of T cell-mediated cross protection than it was observed at the 3-month span. Based on our previous study we believe this protection wanes as macaques exposed to ZIKV 2.8 years after DENV were not afforded this protection. These results suggest that there is a window of optimal T cell cross-protection between ZIKV and DENV.

## Results

### Rhesus macaque cohorts and sample collection

Six rhesus macaques (*Macaca mulatta*) were infected with 5 × 10^5^ pfu s.c. of DENV-2 New Guinea 44 in 2016 (cohort 1 in Fig. 1). In 2017, four rhesus macaques were infected with the same virus strain and pfu (cohort 2 in Fig. 1). In addition, a control group, composed by six flavivirus-naïve rhesus macaques were added as control (cohort 3 in Fig. 1). All three cohorts were infected with 1 × 10^6^ pfu s.c. ZIKV PRVABC59 on the same day, defining exposure time between infections for cohorts 1 and 2 as 12 months or middle convalescent and 3 months or early convalescent, respectively (Fig. 1). Prior to the challenge with ZIKV, all sixteen animals were put through a quarantine period of forty days. Figure 1 also denotes the unexpected setback that Hurricane María brought to our work plan. Sample collection programmed from days 7 to 29 p.i. was interrupted due to inability of access and/or lack of power at the CPRC facilities, University of Puerto Rico, San Juan, Puerto Rico.

**Figure 1.**
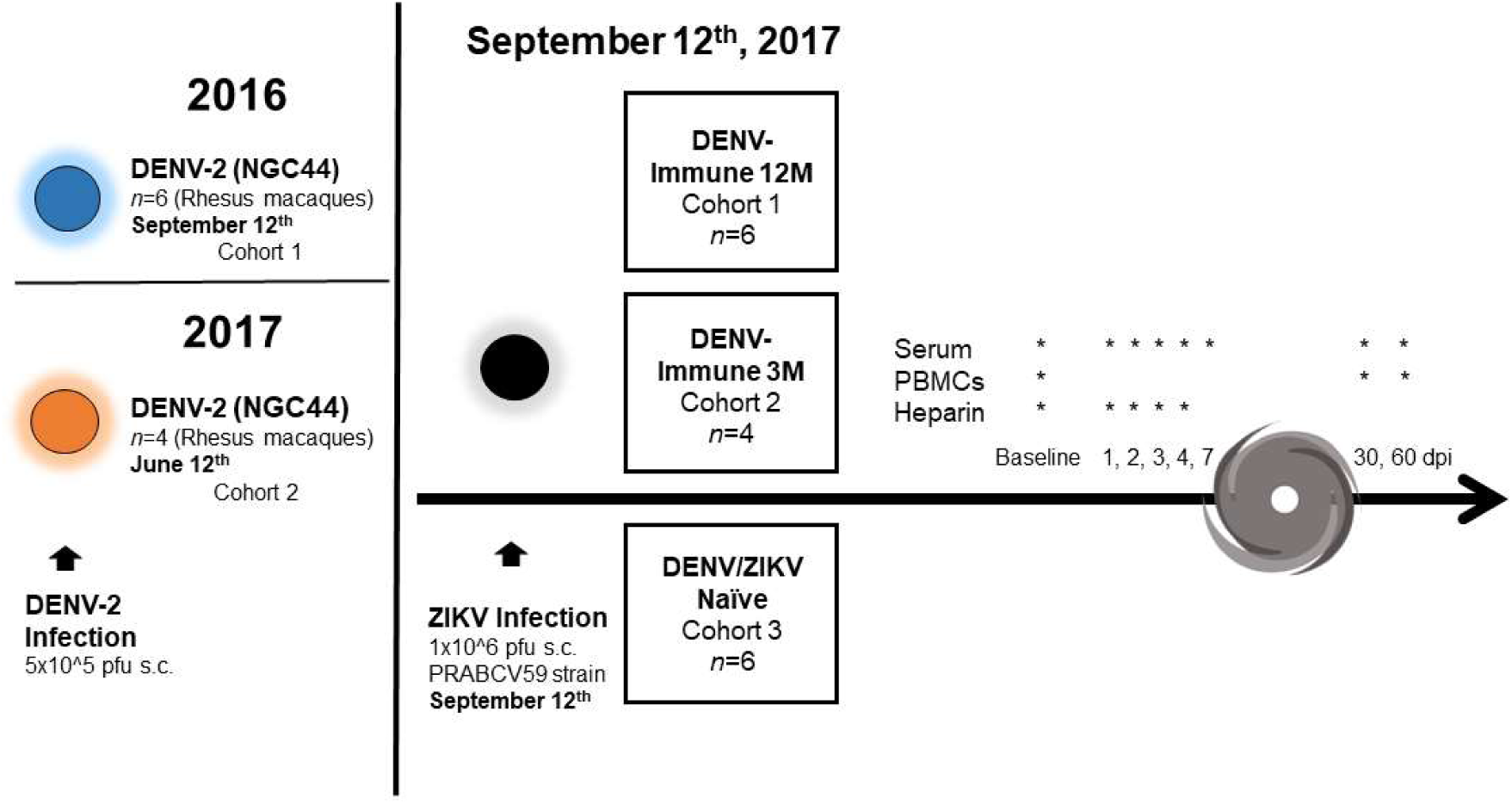
Experimental design of ZIKV infection in DENV-immune and naïve macaques. Two cohorts of rhesus macaques (*Macaca mulatta*) were exposed to DENV-2 (5 × 10^5^ pfu s.c.) at different timepoints. Both cohorts were exposed to ZIKV strain PRABCV59 (1 × 10^6^ pfu s.c.) on September 12^th^, 2017, along with a third cohort composed of zika and dengue naïve animals (n=6). ZIKV infection was performed 12 months after DENV infection for cohort 1 (n=6), and 3 months after DENV infection for cohort 2 (n=4). Serum was collected at baseline and days 1 through 7 post ZIKV infection (p.i.). Sample collection was interrupted by Hurricane María’s impact, and resumed on day 30 p.i. PBMCs could only be obtained on baseline, day 30 and 60 p.i., while heparinized whole blood was collected on baseline and days 1 through 3 p.i. Additionally, urine was collected on baseline and days 2, 4 and 6 p.i.

### Clinical status and laboratory results are affected by DENV immunity

To determine how a previous DENV infection affects the clinical status of non-human primates after a ZIKV infection, day 0 (baseline), day 6 p.i. and day 30 p.i. were compared in terms of complete blood count and liver enzymes levels. All sixteen animals belonging to this study were continuously monitored and evaluated twice daily for evidence of disease or injury. All animals were inside the range in terms of weight (Fig. S1). DENV 3M animals presented a significant increase in external (axillary) temperature compared to DENV 12M and naïve animals at day 5 p.i., followed by a sudden drop by day 6 p.i. (P<0.0005; mean diff.: -2.05, CI95%: -3.31 to -0.80 and P<0.0001; mean diff.: 2.75, CI95%: 1.49 to 4.00, respectively) (Fig 2A). No variations were detected in rectal temperature (Fig. 2B). Liver enzyme aspartate aminotransferase (AST) and alanine aminotransferase (ALT) levels were significantly elevated at day 6 p.i. in the naïve and DENV 12M animals compared to their baseline levels (P<0.001; mean diff.: -23.8, CI95%: -38.15 to - 9.51 and P<0.0001; mean diff.: -33, CI95%: -47.32 to -18.60, respectively), while DENV 3M animals did not present major variations between baseline and day 6 p.i. (Fig. 2C,D). The naïve group had significantly higher values of AST compared to both DENV immune animals (Fig. 2C), and of ALT compared to the DENV 3M group (Fig. 2D) (P<0.05; mean diff.: 18.3, CI95%: 4.01-32.7 and P<0.001 for naïve vs. DENV 12M; mean diff.: 33.2, CI95%: 17.2 to 49.3 for naïve vs. DENV 3M). Values returned to near baseline levels in all three group by day 30 after the infection. These results together suggest that previous immunity to DENV may play a protective role against ZIKV-induced liver damage. All cohorts had a drop in white blood cell counts (WBC) by day 6 p.i. that increased to near baseline levels by day 30 p.i. (Fig. S2A). No significant variations were noticed in platelet counts (PLT) for any of the cohorts (Fig. S2B). Although no differences were detected between groups, monocytes (MON) were significantly higher in the naïve animals by day 6 p.i. compared to their baseline levels (P<0.05; mean diff.: 0.25, CI95%: 0.00 to 0.46) (Fig. S2C,D). This was not observed in the DENV pre-exposed animals.

**Figure 2.**
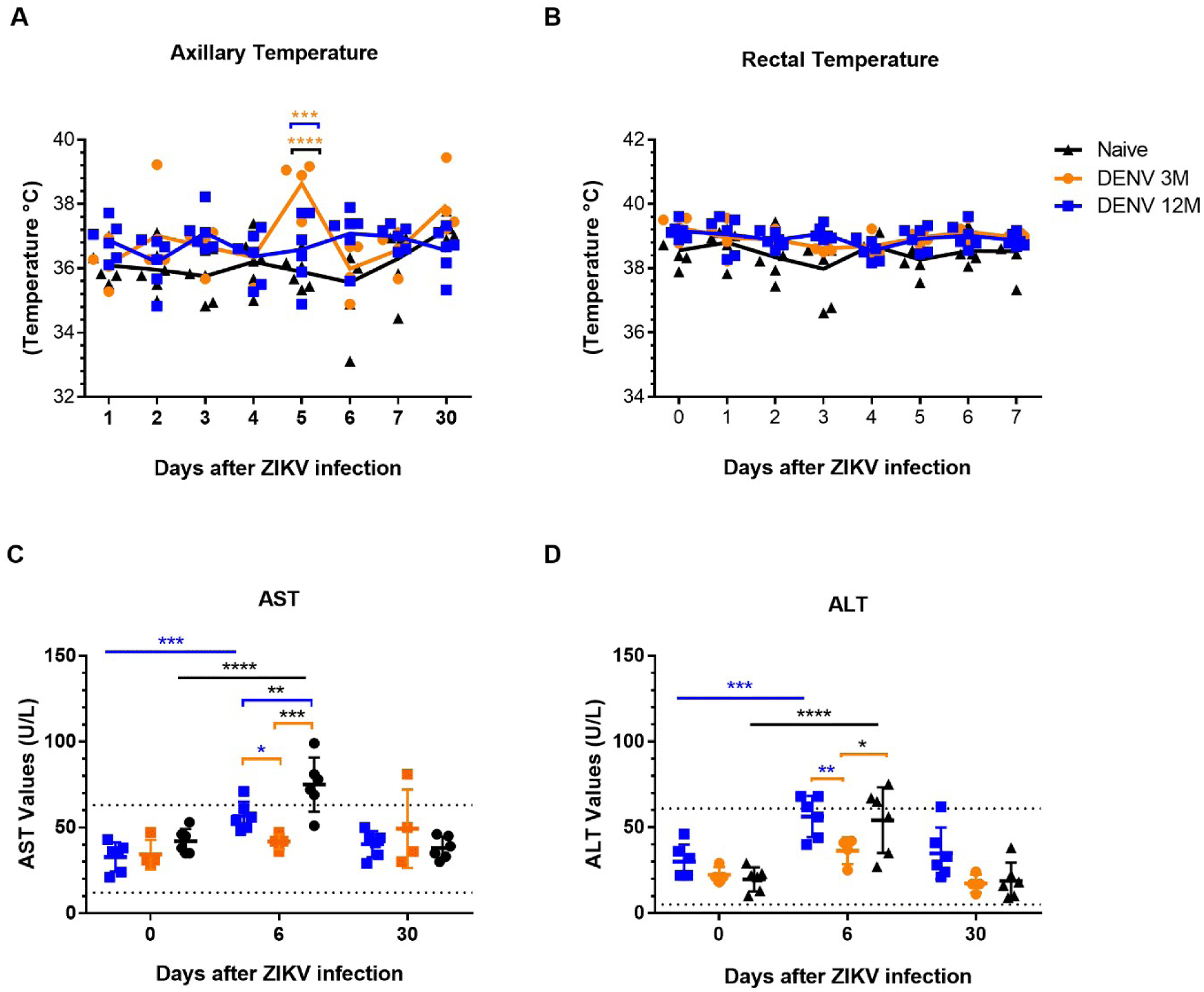
Vital signs and clinical laboratory status of macaques before and after ZIKV infection. Significant changes in the vital signs and laboratory values after ZIKV infection are showed. In all panels, animals exposed to DENV 12 months before ZIKV infection are depicted in blue, while animals exposed to DENV 3 months before are in orange. Naïve animals are in black. (A) External temperature (in Celsius) and (B) rectal temperature (in Celsius) were measured. Statistically significant differences among groups were calculated by two-way ANOVA using Tukey’s multiple comparisons test (***P<0.0005 and ****P<0.0001). (C) Aspartate Aminotransferase (AST) and (D) Alanine Aminotransferase (ALT) levels at different timepoints. Dotted lines represent normal clinical ranges for rhesus macaques. Statistically significant differences among groups were calculated using an unpaired multiple t test, while differences within cohorts in respect with their baseline values were computed by two-way ANOVA using Dunnett’s multiple comparisons test (*P<0.05, **P<0.001, ***P≤0.0001 and ****P<0.0001). Colored stars represent a significantly different group, while colored lines represent the group that it is compared to.

### ZIKV RNAemia is affected by the longevity of previous DENV immunity

To determine if previous immunity to DENV enhances or reduces ZIKV replication, and how it changes depending on the convalescent period, ZIKV RNAemia was measured in serum and urine using qRT-PCR. RNAemia was defined as follows: early from day 1 to 3 p.i., middle from day 4 to 7 p.i., and late viremia from day 7 p.i. onwards (days 30 and 60). During the early period, viral RNA detection increased similarly in all groups (Fig. 3A).

**Figure 3.**
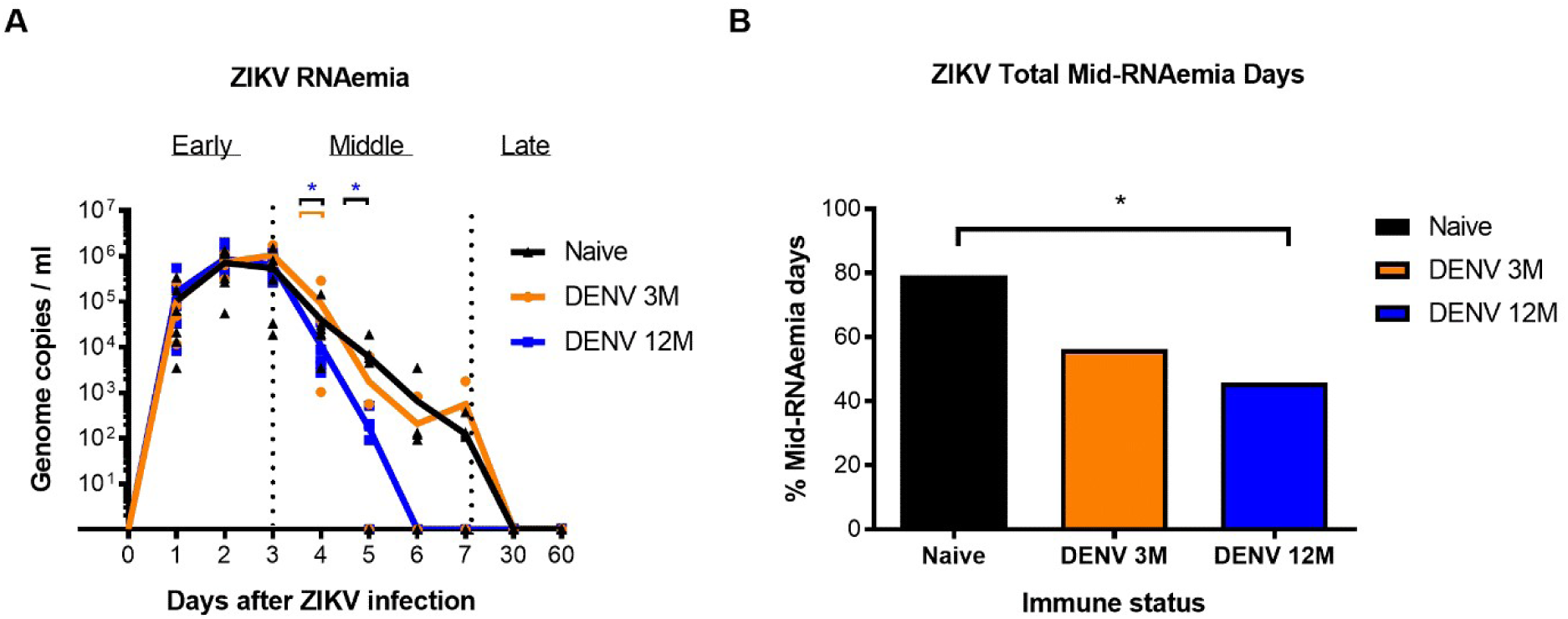
Zika RNA kinetics in serum and RNAemia days per cohort. RNAemia days are affected by convalescence produced after DENV infection depending on time between exposures. In all panels, animals exposed to DENV 12 months before ZIKV infection are in blue, while animals exposed to DENV 3 months before are in orange. Naïve animals are in black. (A) Zika RNAemia was defined as early RNAemia (days 1 to 3 p.i.), mid RNAemia (days 4 to 7 p.i.), and late RNAemia (days 7 p.i. onwards). ZIKV replication was detected in serum during the first 7 days after infection. Statistically significant differences were observed using unpaired multiple t tests (*P<0.05). Genome copies per mL are shown logarithmically. (B) Total mid-RNAemia days were calculated using the following formula: total viremia days divided by total possible viremia days and are expressed as percentage. The obtained values were placed in a contingency table. Statistically significant differences of viremia days were calculated using a two-sided Fisher’s exact test (*P<0.05). Colored stars represent a significantly different group, while colored lines represent the group that it is compared to.

Of note, DENV-middle convalescent animals had significantly lower peak viremia on day 4 compared with the rest of the animals (P<0.042 vs. naïve and P<0.019 vs. DENV3M group). By day 5 all three groups had two animals with undetectable viremia. However, the set-point viremia in the four animals from DENV-12M group was significantly lower compared to the four animals having viremia in the naïve group (P<0.039) (Fig. 3A and Table 1). By days 6 and 7 p.i., there was no viral RNA detection in the DENV-middle convalescent group, while DENV 3M animals showed a trend towards an intermittent viremia and most of the naïve animals still had detectable viral RNA. By days 30 and 60 all animals tested negative for ZIKV (Fig. 3A and Table 1).

**Table 1.**
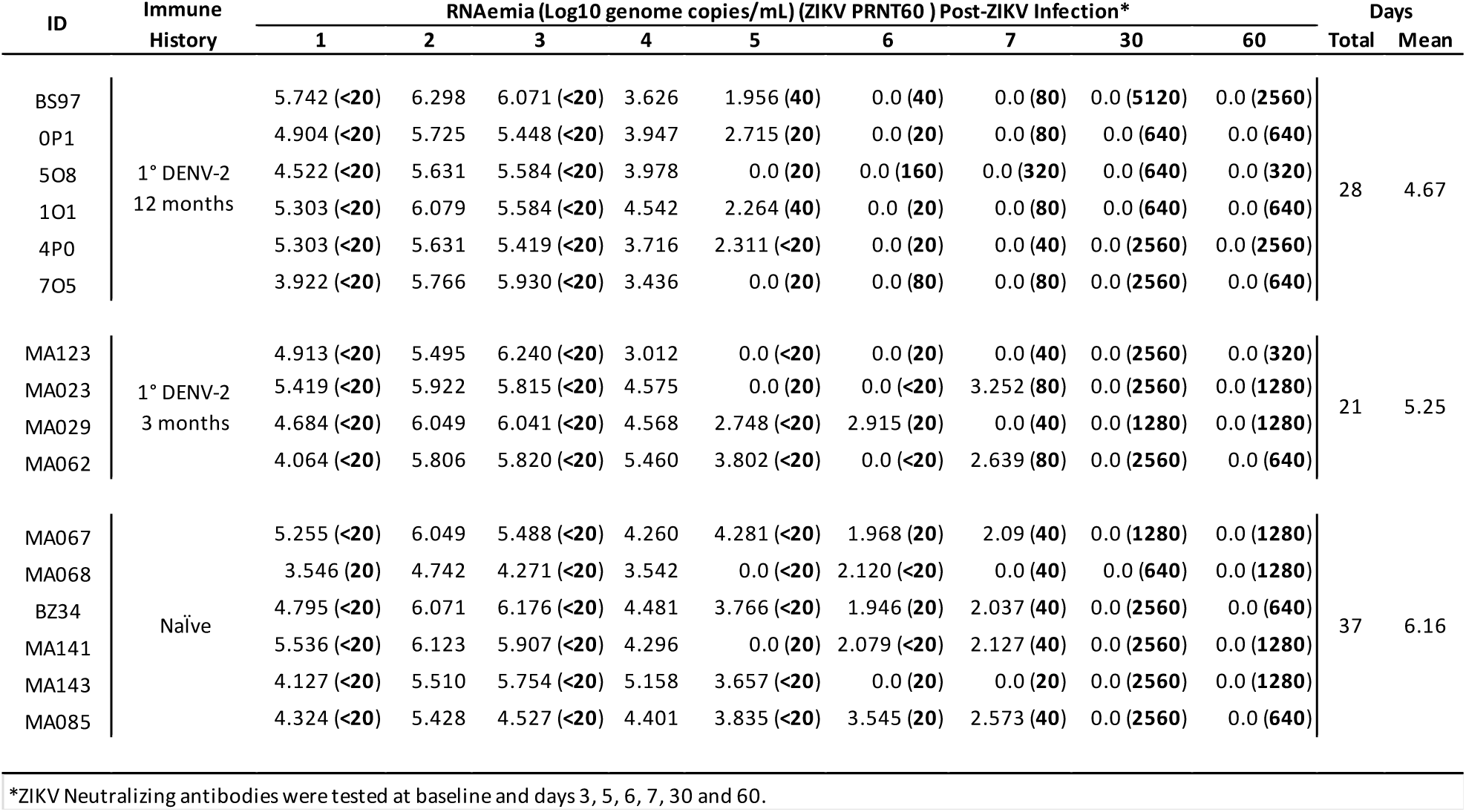
ZIKV RNAemia days of naïve and DENV-immune macaques. ZIKV RNA detection was consistent in all groups during the first 4 days post infection (p.i.). Peak viremia occurred on day 3 p.i. Cohort 1 animals had no detection of ZIKV RNA in serum by day 6 p.i. Mean viremia days per group was calculated using days with detectable RNAemia divided by the number of animals in each group.

We defined total mid-RNAemia days as the days with detectable viremia out of all possible days with detectable viremia during the collection period. The animals exposed to DENV 12 months earlier had the least viremia days in comparison with the naïve group, and the difference was statistically significant (P<0.05) (Fig. 3B). These results suggest that a previous infection with DENV contributes to an earlier and more efficient control of ZIKV viremia in a subsequent infection, but only if at least 12 months (a middle convalescence period) have passed between infections. Lastly, ZIKV vRNA in urine was measured using qRTPCR, but only one animal from the DENV 3M group (MA023) had detectable levels at day 6 p.i. (results not shown).

### Serological profile is modified by the time between the two infections

To assess the impact of previous exposure to DENV at different times in a humoral response against a subsequent ZIKV infection, all sixteen animals were tested for binding antibodies against ZIKV and DENV serotypes following ZIKV infection. All three groups had levels of anti-DENV IgM below the cutoff value during the three collection periods, suggesting ZIKV infection did not induce DENV-specific IgM response (Fig. S3A). As expected, all DENV immune animals had detectable IgG levels against DENV at baseline (Fig. S3B). Anti-DENV IgG levels were confirmed in both DENV-pre-exposed groups and by day 30 experimented a significant expansion compared to their basal levels (P<0.05; mean diff: -0.27, CI95% -0.5145 to-0.02546 P< 0.005; mean diff: -0.3, CI95% -0.4997 to -0.1003 for the DENV3M and DENV12M groups respectively). Those cross-reacting antibodies were also significantly higher compared to the naïve animals on day 30 (P<0.0001; mean diff.: 0.96, CI95%: 0.61 to 1.30 for DENV 12M vs naïve at 30 days p.i,; P<0.0001; mean diff.: 0.75, CI95%: 0.37 to 1.14 for DENV 3M vs naïve at 30 days p.i.; P<0.0001; mean diff.: 0.88, CI95%: 0.54 to 1.23 for DENV 12M vs naïve at 60 days p.i.; P<0.001; mean diff.: 0.68, CI95%: 0.29 to 1.07 for DENV 3M vs naïve at 60 days p.i.), slowly decreasing by day 60 p.i. (Fig. S3B). After a limited increase of the anti-DENV IgG levels on day 30 p.i. the levels rapidly decrease by day 60 in the naïve group. Moreover, by day 30 p.i., DENV-middle convalescent animals showed a strong trend of having higher levels of anti-DENV IgG compared to the DENV-early convalescent animals, although no statistical significance was reached.

As expected, all ZIKV-infected animals developed anti-ZIKV IgM 30 days after the infection (Fig. S3C). However, those antibodies were early detected only in three animals from the DENV12M group with two subjects showing a peak on that day (5O8, 1O1) (Fig. S3C). All DENV-immune animals had detectable levels of anti-ZIKV IgG at baseline compared to the naïve animals, suggesting a strong cross-reactivity between previously generated anti-DENV IgG to ZIKV (Fig. S3D). DENV-early convalescent animals had significantly higher levels of anti-ZIKV IgG than the other DENV-immune animals (DENV12M) at baseline (P<0.05; mean diff.: -0.59, CI95%: - 1.15 to -0.05). By day 7 p.i., DENV-immune animals have higher levels of anti-ZIKV IgG that increase throughout days 30 and 60 p.i.. Similar but slower increase was detected in the naïve group. Nonetheless, all three groups showed a boost in anti-ZIKV IgG levels by day 30 p.i. with a significant expansion at day 60 only in the DENV-immune animals (P<0.0001; mean diff.: - 1.187, CI95%: -1.848 to -0.525 and P<0.0001; mean diff.: -1.176, CI95%: -1.716 to -0.636 for DENV3M and DENV12M groups respectively (Fig. S3D). However, all animals except one in the naïve group also experiment a significant expansion in their level of anti-ZIKV IgG by day 60 compared to day 30 (P<0.0001; mean diff.: -1.231, CI95%: -1.771 to -0.691).

Only one animal from the DENV-early convalescent group (MA062) had very low but detectable anti-ZIKV NS1 IgG levels at baseline (Fig. S3E). All ZIKV-infected animals showed a boost in anti-NS1 levels by day 30 p.i., with DENV-middle convalescent group having significantly higher levels compared to early convalescent and naïve animals (P<0.01; mean diff.: 1.05, CI95%: 0.2608 to 1.836 for DENV12M vs. DENV3M; P<0.0001; mean diff.: 2.15, CI95%: 1.45 to 2.859 for DENV12M vs naïve). These levels decreased by day 60 p.i., and the drop is more dramatic in the pre-immune animals (Fig. S3E). Antibodies against ZIKV EDIII were also measured in order to determine their contribution to humoral immunity and for their known specific contribution to ZIKV neutralization 25(Fig. S3F). Only one animal from the DENV-middle convalescent group showed detectable levels of anti-ZIKV EDIII before ZIKV infection. Anti-ZIKV EDIII levels for all groups slowly increased throughout 30 and 60 days p.i., and the increase at day 60 p.i. was significant for DENV-immune animals with respect to their basal levels (P<0.05; mean diff.: -0.85, CI95%: -1.68 to -0.03 for DENV12M animals, and P<0.001; mean diff.: -1.26, CI95%: -2.27 to -0.25 for DENV3M animals). No significant differences were observed among the groups, suggesting that previous exposure to DENV does not have an impact on the generation of cross-reactive antibodies against ZIKV EDIII epitopes and that these antibodies may have limited contribution to ZIKV neutralization.

### The time between DENV/ZIKV infections modifies the neutralizing profile

To determine the contribution of binding antibodies to the neutralization and the impact of a previous DENV infection in a subsequent ZIKV infection in terms of neutralization potential, all three groups were tested using PRNT and FRNT assays against ZIKV and the four DENV serotypes respectively. Neutralization assays were completed for baseline, days 30 and 60 for DENV and ZIKV. To better understand if the previous DENV immunity plays a role in the neutralization of ZIKV, we also ran PRNT assay for days 3, 5, 6 and 7. The endpoint titers are showed in Table 1 along with the viremia set point values in order to facilitate a better interpretation of the relationship between both parameters.

In figure 4 the 50% effective concentration (EC) of neutralizing antibodies is shown. As expected, all three groups had low or absent neutralizing antibodies against ZIKV at baseline (Fig. 4A). As early as 6 days after the infection an increase in the neutralizing activity against ZIKV is detected in all groups with a slight non-significant trend to be higher in the DENV12M group. This increase continues on day 7 with the trend to be higher in both preimmunized groups. By day 30 p.i., neutralizing titers had boosted in the three groups, with DENV-middle convalescent animals showing a slight trend towards higher levels of dilution effective for half-maximum neutralization compared to DENV3M and naïve groups. These levels decline slightly by day 60 p.i but still maintained a similar relation among groups (Fig. 4A). In order to expand our analysis of the contribution of the humoral immune response to the early viral replication we further looked at the dilution:neutralization capacity relation in samples from all timepoints. As shown in figure 4C, we identified significant differences, at the highest serum concentrations in the magnitude of the neutralization (only dilutions showing more than 60% of neutralization were considered) on days 6 (P<0.001 and P<0.029 for DENV12M and DENV3M vs. naïve respectively at 1:20 dilution and P=0.0005 and P<0.0001 for DENV12M vs. DENV3M and naïve respectively at 1:40 dilution) and 7 (P<0.0039 and P<0.0001 for DENV-12M vs. naïve at 1:20 and 1:40 dilutions respectively) p.i.. However, the role of the neutralization limiting viral replication, particularly at day 6 after the infection, is debatable when the relationship between RNAemia and end point neutralizing titers are analyzed together (Table 1).

**Figure 4.**
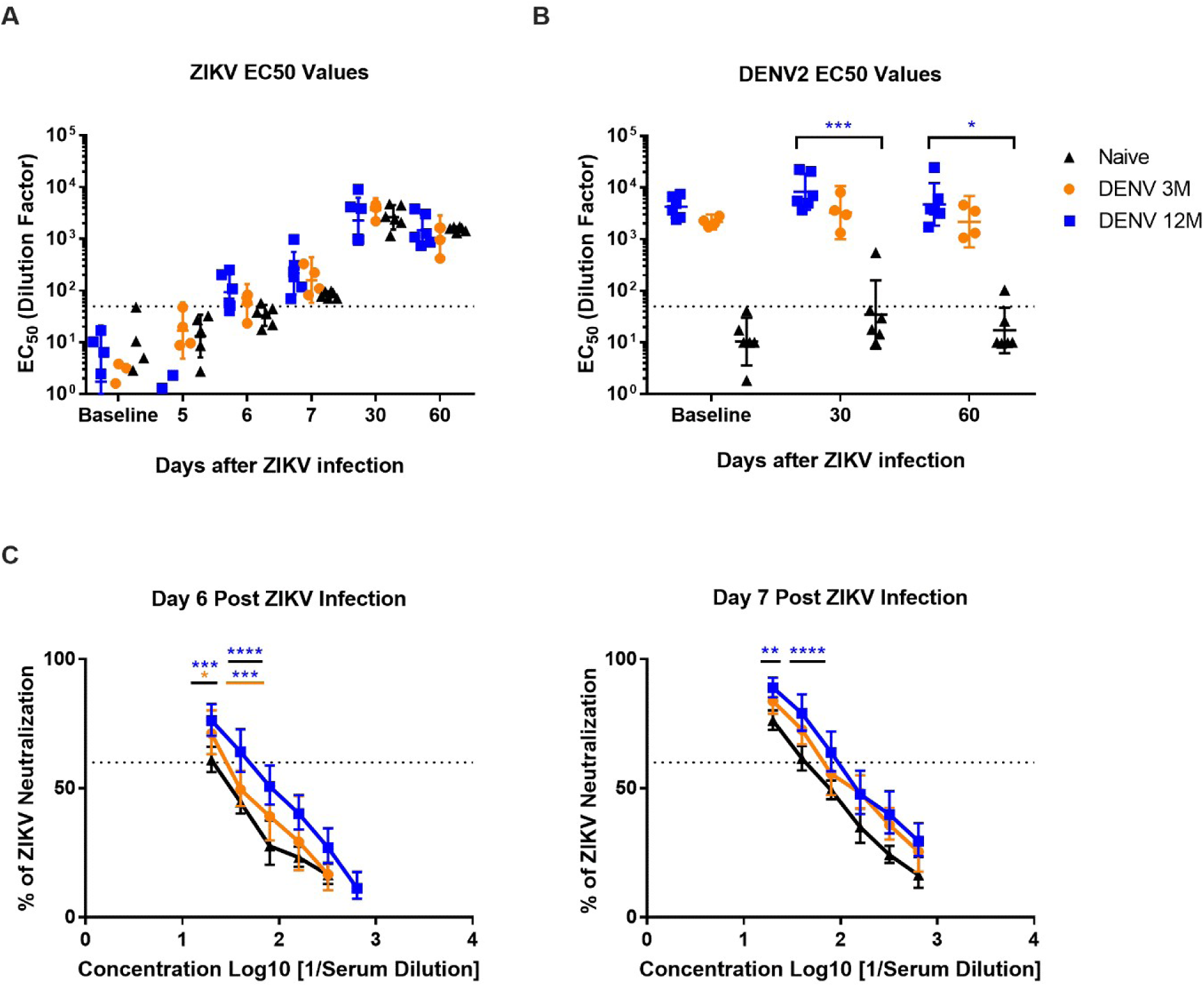
Geometric mean titers of dengue and ZIKV neutralizing antibodies. The 50% effective concentration of neutralizing antibodies was determined. Animals from cohort 1 are shown in blue, animals from cohort 2 are shown in orange and naïve animals from cohort 3 are shown in black in all panels. Dotted line indicates the limit of detection for the assay. Non-neutralizing sera were assigned a value of one-half of the limit of detection for visualization and calculation of the geometric means and confidence intervals. (A) EC50 values of neutralizing antibodies against ZIKV after ZIKV infection. (B) EC50 values of neutralizing antibodies against DENV2 after ZIKV infection. Statistically significant differences among groups were calculated by two-way ANOVA using Tukey’s multiple comparisons test (*P<0.05 and ***P≤0.001). (C) Dilution titers against ZIKV are shown during day 6 and 7 post ZIKV infection. Statistically significant differences among groups were calculated by two-way ANOVA using Tukey’s multiple comparisons test (*P<0.05, **P<0.001, ***P≤0.001 and ****P<0.0001). Colored stars represent a significantly different group, while colored lines represent the group that it is compared to.

Both DENV immune groups had high levels of neutralizing antibodies against DENV2 at baseline, which boosted significantly for DENV12M animals at day 30 p.i. compared to the naïve group (Fig. 4B). By day 60 p.i., these neutralizing antibodies did not decline in neither of the DENV immune groups, and a significant difference was still present for DENV-middle convalescent animals, which suggests that ZIKV infection induced a boost in cross-neutralizing antibodies to DENV and the magnitude of the boost depend on the time elapse between DENV and ZIKV infection (P=0.0006 for day 30 p.i. and P=0.02 for day 60 p.i.). Only one naïve animal produced low level of neutralizing antibodies against DENV2 by day 30 p.i. that declined by day 60 p.i. (Fig. 4B). Using the anti-ZIKV IgG data presented on the previous section (Fig. S3D), we can conclude that the expansion 60 days p.i. is supported in cross-reactive non-neutralizing antibodies (Fig. 4). Detailed dilution end points for each animal are shown in Figure S4, panels A and B.

When evaluating the neutralizing titers against the all four DENV serotypes, we observed a boost in neutralization against all serotypes in all three groups, suggesting that a subsequent ZIKV infection impact the levels of heterologous DENV-neutralizing antibodies (Fig. S5A,B). Interestingly 30 days after ZIKV infection there was a non-significant trend to higher neutralizing titers against DENV2 and DENV4 compared to the other two DENV serotypes in the DENV naive group. The hierarchy of neutralizing antibodies generated 30 days p.i. was the same for both DENV immune groups (D2>ZIKV>D4>D3>D1), and for the naïve group it was ZIKV>D4>D2>D3>D1 (Fig. S5B). In order to determine if there were any strain-specific neutralization differences, neutralization assays were performed at 30 days p.i. against two recently circulating contemporary ZIKV strains, ZIKVH/PF/2013 and ZIKVPRVABC-59. As showed in figure S5C, no differences in the neutralization magnitude were seen for any group. Altogether, these results confirm the contribution of pre-existing DENV immunity to the expansion of cross-reactive anti-ZIKV IgG levels and of the DENV cross-neutralizing antibodies early after ZIKV infection in macaques.

### Immune cell subsets frequency is shaped by previous DENV exposure

3To establish how previous immunity to DENV shapes the cellular response against a subsequent ZIKV infection, an analysis of the involved cells was performed. Animals exposed to DENV three months earlier had significantly higher frequency of B cells (CD20+) 24 hours before ZIKV infection compared to the other groups (P<0.05; mean diff.: -12.33, CI95%: -22.69 to -1.979 for DENV3M versus DENV12M, and P<0.01; mean diff.: 14.77, CI95%: 4.41 to 25.12 for DENV3M versus naïve group). No other differences between groups were detected, although the trend observed in day 0 is maintained through day 3 (Fig. S6A). On the other hand, the frequency of activated B cells (CD20+CD69+) was very similar in all three groups (Fig. S6B). In addition, we characterized the CD4+ and CD8+ T cells central memory (TCM) and effector memory (TEM) subsets in order to determine how a previous DENV infection impacts the differentiation to these compartments. The frequency of TCM CD4+ cells (CD4+CD3+CD28+CD95+) was significantly lower for the DENV-middle convalescent group compared to the DENV-early convalescent and naive groups at baseline (P<0.05; mean diff.: -19.39, CI95%: -33.23 to -5.35 versus DENV 3M and P<0.05; mean diff.: -19.5, CI95%: -31.97 to -7.03 versus naïve), day 1 p.i. (P<0.05; mean diff.: -14.53, CI95%: -28.47 to -0.59 versus DENV 3M and P<0.05; mean diff.: -1.77, CI95%: -14.23 to 10.7 versus naïve) and day 3 p.i. (P<0.05; mean diff.: -22.97, CI95%: -36.9 to -9.03 versus DENV 3M, and P<0.05; mean diff.: -17.22, CI95%: -29.68 to -4.75 versus naive) (Fig. S7A). On the other hand, the frequency of TEM CD4+ cells (CD4+CD3+CD28-CD95+) was increased before ZIKV infection and remained steady throughout the four timepoints for the DENV-middle convalescent animals, in comparison with the other two groups. Referring to the CD8+ TCM and TEM cells, a similar trend can be observed, although it only reaches statistical significance at day 3 p.i. for TCM cells (P<0.05; mean diff.: -9.85, CI95%: -17.02 to -2.68 versus naïve animals), and at day 1 p.i. for TEM cells (P<0.05; mean diff.: 19.57, CI95%: 0.09 to 39.04 versus DENV 3M animals) (Fig. S7B). Additionally, naïve animals had significantly higher levels of TCM CD8+ cells compared to the DENV immune groups at baseline (P<0.05; mean diff.: - 15.47, CI95%: -22.63 to -8.30).

We noticed a trend that is especially prominent in DENV-middle convalescent animals, of higher TEM cell and lower TCM cell frequencies. In order to scrutinize this pattern, we analyzed the pattern of frequency in each group separately (Fig. S7C,D). Compared to their baseline values, only naïve animals had significant differences on days 1 through 3 p.i. between CD4+ and CD8+ TEM and TCM subsets (P<0.05). Additionally, the naïve animals go through a sudden contraction of TCM CD4+ and CD8+ cells at day 1 p.i. that slowly begins increasing by day 3 p.i. (P<0.05). Similarly, the TEM CD4+ and CD8+ cells expand, although TCM cells reach parallel levels by day 3 p.i. (P<0.05). This phenomenon can be seen in DENV 3M animals in the TCM CD8+ cells during day 1 p.i., but not TCM CD4+ cells. DENV 12M animals go through the same occurrence, but reaching statistical difference in all three days p.i (P<0.05) (Fig. S7C,D). These results are confirmed by the pattern showed by individual animals showing that DENV-middle convalescent animals had a more pronounced TEM cell expansion and TCM cell contraction compared to their DENV-early convalescent counterparts (Fig. S8). Noticeable, this contraction of TCM cells and expansion of TEM CD4+ T cells, while it is not antigen specific, translate into a more efficient viremia control in the 12M group but not in the naïve or DENV 3M groups, (Fig 5A,B). Following this, we wanted to compare the level of activation and proliferation of those cell subsets. We found that DENV-early convalescent group had significantly lower proliferation levels of CD4+ TCM cells at day 2 p.i. compared to the other two groups (Fig. S9A) and there were no other significant variations observed in terms of proliferation (Fig. S9A-D). In contrast, same group of animals showed a trend to higher activation levels of CD4+ and CD8+ TCM and TEM cells in comparison with the other two groups, although no statistical differences were found (Fig. S9E-H). These findings suggest that the time lapse between DENV and ZIKV infections shapes the cellular immune response against ZIKV.

**Figure 5.**
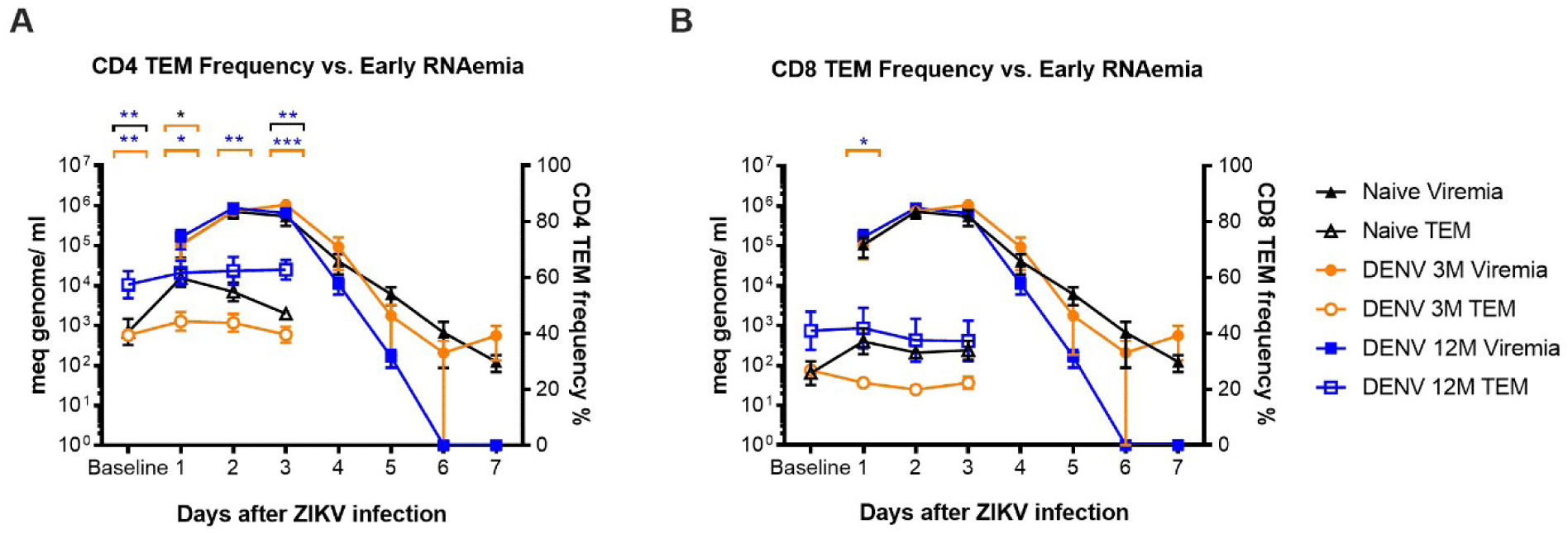
CD4 but not CD8 effector memory cell frequency is associated to early decreased ZIKV RNAemia. Effector memory T cell frequency up to day 3 p.i. is shown in comparison to RNAemia (days 1 to 7 p.i.). Animals exposed to DENV 12 months before ZIKV infection are in blue, animals exposed to DENV 3 months before are in orange and naïve animals are in black. (A-B) Genome copies per mL are shown logarithmically, represented by filled colored symbols (left Y axis), while CD4 and CD8 effector memory T cell frequency is represented by hollow symbols (right Y axis). Comparisons between cohorts were performed by two-way ANOVA using Tukey’s multiple comparisons test (*P<0.05, **P<0.001 and ***P≤0.001). Significant statistical differences shown belong to effector memory T cell frequency. Colored stars represent a significantly different group, while colored lines represent the group that it is compared to.

### T cell immune response is boosted by the time lapse between DENV/ZIKV infections

To assess if a previous DENV infection has an impact on the T cell response to a ZIKV infection, their effector responses were measured. CD4+ and CD8+ T cells produced IFNg, TNFa, and CD107a in response to various stimuli (Fig. 6A-B). The IFNg response in the CD4+ T cells from the DENV 12M group before ZIKV infection is remarkable. The frequency of these cells was significantly higher in response to the whole inactivated DENV (P<0.05) and showed a strong trend to have a higher frequency of IFNg producing cells in response to peptides derived from the DENV and ZIKV envelopes and ZIKV non-structural proteins as well compared to the 3M and naïve groups.

**Figure 6.**
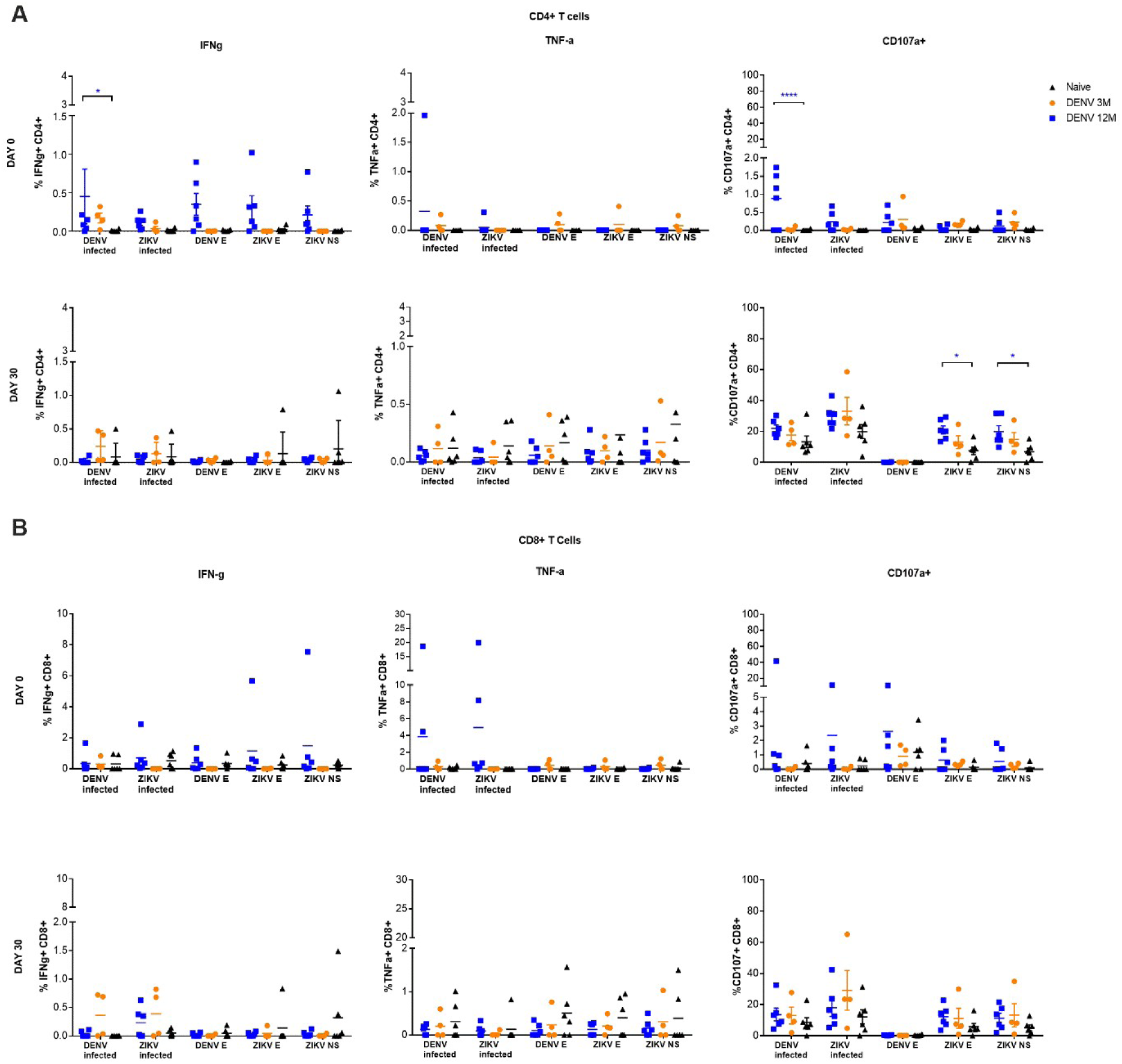
Antigen-specific CD4^+^ and CD8^+^ response prior and after ZIKV infection. The frequency of the specific response to DENV and ZIKV antigens differs among cohorts. In all panels, animals exposed to DENV 12 months before ZIKV infection are in blue, while animals exposed to DENV 3 months before are in orange. Naïve animals are in black. All percentages shown are subtracted from the unstimulated background. (A) Analysis of CD4 T cell response to different stimuli before (upper panel) and 30 days after ZIKV infection (lower panel). (B) Analysis of CD8 T cell response to different stimuli before (upper panel) and 30 days after ZIKV infection (lower panel). Statistically significant differences among groups were calculated by two-way ANOVA using Dunnett’s multiple comparisons test (*P<0.05 and ****P<0.0001). Colored stars represent a significantly different group, while colored lines represent the group that it is compared to.

DENV 12M animals had also a significantly higher frequency of CD107a+ cells prior to ZIKV infection (P<0.0001; mean diff.: -0.8775, CI95%: -1.25 to -0.503), while a significant increase in reactivity of CD107a+ CD4+ cells was observed against ZIKV envelope and non-structural antigens 30 days p.i. (P<0.05; mean diff.: -0.4421, CI95%: -0.8616 to -0.02263 and P<0.05; mean diff.: -0.8775, CI95%: -1.251 to -0.5035, respectively) (Fig. 6A). These results correlate with the protective effect observed in this group (Fig. 6A). Nothing remarkable was observed in TNFa CD4+ frequency. In contrast, data from CD8+ T cells denote similar responses between DENV immune animals, with no significant variations compared to the naïve animals (Fig. 6B). This suggests that previous DENV immune status preferentially shapes the CD4+ T cells effector responses to a ZIKV infection. Gating strategy is provided as supplementary figure 10.

### Pro-inflammatory cytokines are not exacerbated by previous DENV immunity

Next we determined how DENV immunity impacts the cytokine secretion during a subsequent ZIKV infection (Fig. 7). Naïve macaques had significantly higher levels of pro-inflammatory cytokine IFN-a on day 1 p.i. compared to the other groups (P<0.05; mean diff.: -145.6, CI95%: -271.7 to - 19.37) (Fig. 7A). This trend continued through the rest of the collection period, but no other significant differences were detected. DENV 12M animals had seemingly higher levels compared to their DENV 3M counterparts, although no statistical significance was reached. A similar event can be observed with CXCL10, where naïve animals had a significant boost by day 1 p.i. in comparison with the other two DENV-immune groups, although a dramatic drop occurs on the following 24 hrs (P<0.05; mean diff.: -533, CI95%: -22.69 to -1.98) (Fig. 7B). On day 2 the levels of CXCL10 increase prominently on the DENV immune animals, noticeably higher and more consistent on the DENV 12M group, but no significant differences are noted. This same group of DENV-middle convalescent animals showed a trend towards higher MIP-1a levels, reaching significant differences in day 5 and 7 p.i. (P<0.01), while naïve animals showed an increase in MIP-1b levels by day 1 p.i. (P<0.01), followed by a sudden drop (Fig. 7C and D, respectively). Likewise, an increase in IL-1Ra levels, which is considered an inflammatory marker, was detected in naïve animals at day 1 p.i. that is significantly higher than levels in DENV-immune animals (P<0.01; mean diff.: -1723, CI95%: -3060 to -385.1 versus DENV 12M; P<0.01; mean diff.: -1809, CI95%: -3305 to -313.18 versus DENV 3M), but it decreases in the next 24 hours (Fig. 7E). On the other hand, a significant increase in BAFF levels at day 5 p.i. was observed in DENV 12M animals (P<0.05; mean diff.: -3044, CI95%: -5999 to -87.73) (Fig. 7F). Interestingly, animals exposed to DENV 12 months before ZIKV infection showed higher levels of circulating perforin, reaching a significant difference compared to the other two groups at day 7 p.i. (P<0.05; mean diff.: -419.9, CI95%: -824.2 to -15.68) (Fig. 7G). This result supports a role for perforin cytotoxicity in early viral clearance.

**Figure 7.**
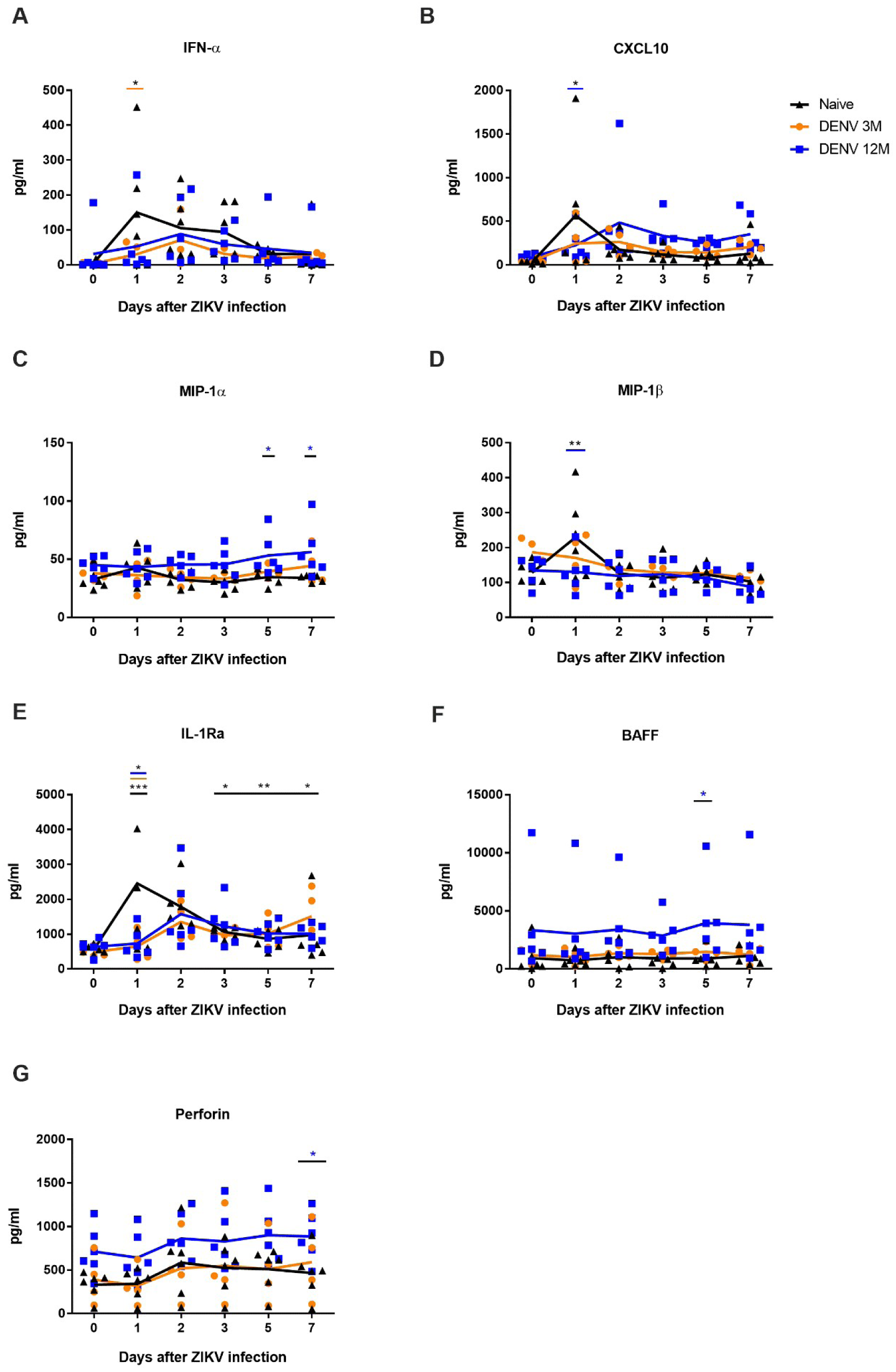
Previous exposure to DENV modulates the cytokine and chemokine profiles after ZIKV infection. (A-G) Significant cytokine and chemokine profiles of are depicted in pg per mL. In all panels, animals exposed to DENV 12 months before ZIKV infection are in blue, while animals exposed to DENV 3 months before are in orange. Naïve animals are in black. Statistically significant differences among groups were calculated by two-way ANOVA using Tukey’s, Sidak’s and Dunnett’s multiple comparisons tests (*P<0.05, **P<0.001 and ***P≤0.0001). Colored stars represent a significantly different group, while colored lines represent the group that it is compared to.

### Other immune cell responses

Plasmacytoid dendritic cell (pDCs) frequency could not be measured on baseline and day 1 and 2 p.i. due to a staining problem. Nonetheless, day 3 p.i. data presents no detectable differences in the frequency of dendritic cells of pDC lineage or absolute pDCs between groups (results not shown). In contrast, dendritic cells of myelocytoid lineage (mDCs) show a significant increase in frequency at days 1 and 2 p.i. in DENV 12M animals (P<0.01; mean diff.: 10.54, CI95%: -2.65 to 18.43 for DENV 12M versus naïve animals on day 1 p.i.; for day 2 p.i., P<0.01; mean diff.: 13.68, CI95%: 4.87 to 22.50 for DENV 12M versus DENV 3M and, P<0.0001; mean diff.: 15.57, CI95%: 7.68 to 23.45 for DENV 12M versus naïve animals)(Fig. S11). This suggests that an early activation of mDCs induced by previous immunity to DENV correlates with ZIKV control. This result is coherent with previous finding that CXCL10 (Fig. 7B) frequency was higher in DENV immune animals after day 2 p.i., especially in those exposed to DENV 12 months before, supporting control of viremia in this group.

## Discussion

It is well established that exposure to DENV prior to a ZIKV infection results in a qualitative modification of the humoral and cellular immune response to ZIKV in mice, macaques and humans (24, 26-29). However, we know from early human temporal challenge studies with multiple serotypes of DENV that the timing between flavivirus exposure can alter the balance between heterologous immune pathogenesis and protection. In human populations living in flavivirus endemic areas, it is difficult to establish the exact time of a primary infection and more difficult to establish the time between two consecutive infections.

Macaques provide a robust model to study the immunological profile after sequential flavivirus infections (24, 30, 31). Using the macaque model allows among other factors, to normalize the quantity of viral inoculum, the age, and sex of the animals exposed, and the timing of the infection. Controlling the timing of exposure between heterologous flavivirus infections allows for the measurement of the impact of time on the immune response to the second infecting agent. These studies are important not only for understanding the impact duration of exposure of heterologous natural flavivirus infections in endemic areas but also provides a model to evaluate possible pan-flavivirus protective windows for vaccine candidates.

Previously we have shown that a DENV infection 2.8 years (late convalescence period) prior did not lead to an enhanced ZIKV infection. Moreover, that period of convalescence results in an immune status that trends toward the control of ZIKV viremia and the decrease of liver enzymes after the infection. Differences were also observed in B-cells and T-cells activation and cytokines and chemokines profile (24). Our previous results have been recently validated in a metanalysis using most of the NHP available data (32) and more relevant, in two studies presenting real settings of humans living in DENV-endemic areas (32-34). In this work, we aimed to establish the contribution of different DENV convalescent periods (3 months or early and 12 months or middle convalescent periods) on the immune response and disease course after ZIKV infection. From our results, we confirmed that the length of time after DENV infection significantly impacts ZIKV infection and pathogenesis by limiting an increase in body temperature, controlling viremia, and mitigating liver-induced damage.

It is well documented that secondary flavivirus infections, including ZIKV, lead to an increase in cross-reactive Abs and nAbs ^(28, 35-37)^. We found a transient but significant expansion in the magnitude of the DENV cross-reactive Abs in the pre-immune groups 30 days after ZIKV infection with a rapid decline by day 60. We also confirmed the preexistence of cross-reacting but non-neutralizing antibodies to ZIKV in the DENV-immune groups (35, 36). However, those antibodies were significantly higher only in the DENV-early convalescent group compared to the middle convalescent group. This confirms the high frequency of ZIKV cross-reacting antibodies during the early DENV convalescence that wane during the middle and late convalescent periods (35, 37). Similarly, the levels of the antibodies to NS1 was significantly higher in the group with a middle convalescent period to DENV compared to the other two groups suggesting the presence of more mature memory B cells in this group in comparison with the other DENV immune group. Cumulatively from our previous results(24) and this current work, we can conclude that the neutralization against ZIKV is very limited or absent in all samples we tested after DENV and before ZIKV infections, regardless of the convalescence status. Our results are in agreement with recent findings by Montoya et al. (37) and in partial agreement with other reports using human samples (35, 38-40). The discordance may be related to the differences in the definition of the convalescence period in those previous works.

Taking advantage of our model we completed a detailed analysis between the nAbs and the RNAemia within the first week of ZIKV infection that otherwise would be difficult to conduct in humans’ cohorts. While we identified a significant decline in the viremia set-point on day four in the DENV-12M group only, starting from day 5 to 7 we characterized a limited increase in the magnitude of ZIKV neutralization activity in the DENV12M group. That early increase was characterized by the presence of low-to-intermediate levels of neutralizing antibodies but no correlation can be made with the viremia on days 5 and 6. However, by day 7, only in the animals with DENV-middle convalescence immunity a consistent increase in the ZIKV neutralizing antibodies correlating with the absence of RNAemia was observed only in the animals with DENV-middle convalescent immunity. This result suggests that the humoral immune response in subjects with previous DENV-immunity may contribute to controlling ZIKV replication around one week after the infection. However, the time elapsed between the DENV and ZIKV exposures seems to be a critical factor for that contribution. This result contrast we results from our group suggesting that low-to-intermediary levels of cross-NAbs against DENV induced by a previous ZIKV infection may play a role in controlling the early DENV RNAemia set point (41).

As we discuss below, other mechanisms, most likely associated with the preexisting DENV-induced cellular immune response may be relevant providing the initial control of ZIKV replication. Our findings on the neutralizing activity against ZIKV in the presence of DENV immunity are in agreement with previous works with human mAbs confirming an early expansion of the plasmablast response 6 days after secondary DENV (42, 43) and primary ZIKV infection (44). Particularly interesting is our finding in the DENV12M group characterized by the increase in ZIKV neutralization activity and an early peak of IgM by day 7 in some animals. Previously Lai. et al confirmed that the early expansion of the antibody-secreting B cells secrets ZIKV-specific immunoglobulins G, A but also M (45). Moreover, germinal centers (GC) are responsible for generating long-lived, high-affinity Antibody Secreting Cells (ASC) that in turn are able to generate a better-quality response against re-infections (46). Interestingly, our results show a significant increase in BAFF level (a key factor in germinal center maintenance, B cell maturation, and antibody production) at day five for the animals exposed to DENV 12 months earlier, suggesting a GC response involvement in this group.

This new information fills a current gap in our understanding of the early immune response to ZIKV in the presence of previous DENV immunity and have enormous implications for the diagnostic interpretation and the epidemiological considerations during a flavivirus epidemic and to properly dissect the immune response to ZIKV. A significant body of research has been published so far and seminal conclusions have been drawn without considering the length of time between primary and secondary flavivirus infection. The above-discussed results provide novel insights into the dynamic of the antibodies response to ZIKV in a population previously exposed to DENV.

One of the most important questions still remaining is the role of time between DENV and ZIKV exposure in the protective capacity of the memory T and B cell responses over time. As the time between ZIKV and DENV exposure can vary from months to years, we used our established NHP model of heterologous flavivirus exposure (24) to understand the phenotype of T cells and their protective capacity following heterologous flavivirus infection. Uniquely, we determined that the magnitude and the breadth of the cellular immune response were dependent on the convalescent status with significant consequences. Our study suggests that TEM cells may play a role in controlling ZIKV early set point viremia and as consequence contributing to mitigate the liver damage before the neutralizing antibodies may be effective after a period of mid convalescence, defined on this work as 12 months. That group showed a significantly higher frequency of pre-existing TEM CD4+CD3+CD28-CD95+ cells prior to and continue expanding until day 3 (the last time point tested) after ZIKV infection. Particularly relevant is the significant preexistent combination of high TEM and lower TCM cell frequencies in this group exposed to DENV 12 month prior to ZIKV infection. One caveat of this analysis is that the increase in the frequency of the CD4+ TEM cells observed was not antigen-specific. But the increase from baseline to day 3 was sustained and expanded after ZIKV infection and that expansion correlates with the significant ZIKV set point viremia decrease starting on day 5 after the infection. Protective memory is thought to be mediated by TEM cells that migrate to inflamed peripheral tissues and display immediate effector functions, whereas T central memory (TCM) cells have limited effector function early but have maintained their proliferative capacity (47). Circulating TCM and TEM cell populations can be detected up to 10 years after antigenic stimulation, and their presence correlates with protection as their frequencies increase following booster immunization (48).

A limited but statistically significant proliferation of the CD4+ TCM cells was noted two days after the infection only in the naïve and 12M DENV-immune groups compared to the 3M DENV-immune group. However, proliferating CD4+ and CD8+ T-cells have been reported by 6–8 days post-ZIKV infection (30, 49, 50). Unfortunately, as a consequence of the devastating impact of hurricane Maria, we were unable to collect samples from day 7 until day 30 p.i. in our cohorts. However, from the time points we were able to analyze after ZIKV infection we detected a contraction of the TCM and an expansion of the TEM respectively in naive animals for both the CD4+ and CD8+ T cells compartments, strongly suggesting that the changes in magnitude in those types of cells were specific for ZIKV infection. Of note, the expansion and the contraction of the CD4+ TEM and TCM respectively, were significant compared to their baseline values only in the naïve groups but it was insufficient for effective control ZIKV viremia or to limit the hepatic insult.

Worth to mention, the number of the memory T cells in our model, in spite of the shift from one phenotype to other (TCM>TEM), remains relatively constant over the time in all groups, which is consistent with the proposed mechanism of T cell memory homeostasis (51).

In this study, we are able to combine our previous insights into the role of TCM and TEM cells on prior heterologous flavivirus infection (24) to our current work on the temporal boundaries of immune protection from a heterologous flavivirus infection. From our work, the role of the cellular immune response in facilitating the initial significant decrease of ZIKV replication between days 4 to 7 is very likely. The group challenged with ZIKV 12 months post-DENV infection was better at controlling ZIKV viremia, together with lower levels of the liver enzyme AST compared to the naïve group.

This time-dependent effect has been previously well characterized for the humoral immune response to secondary DENV infections (15, 17, 52). While the transition from central to effector and memory phenotype is a complex and progressive process, the limited response observed in the DENV-early convalescence group may be related to a possible ongoing period of T cells contraction (53) after the clearance of the primary DENV occurs only 3 months earlier. Boosting with a homologous alphavirus replicon before the cell’s contraction period was completed, did not further increase the T cell response in a mice model (54). After YFV and vaccinia virus vaccination in humans, the period of time of T cells contraction is still ongoing around 84 days after vaccination (55). That period of time is similar to the 90 days for the secondary challenge with ZIKV in the DENV3M group. The presence of a mature immune response as a consequence of a previous stimulus 12 months earlier, may explain the contribution of the CD4+ T cells detected in the DENV-middle convalescence group that otherwise is not present in the DENV3M group with limited immune response capabilities or in the naïve group in response to a primary viral infection.

In addition, the cytotoxic profile of the CD4+ T cells present 12 months after DENV infection and during heterologous ZIKV challenge correlate with better performance relative to the early (3 months) or late (2.5 years) (24) periods of time after the primary DENV infection. The role of CD4+T cells in flavivirus infection has been extensively documented (56) (57). Importantly, Weiskopf et al. and others have also shown that DENV CD4+T cells are readily detectable early following DENV infection, and the frequency of DENV-specific CD107a+ CD4+T correlate with enhanced protection against DENV disease (58, 59) and play a key role in controlling secondary flavivirus infections (26). Our work builds on these observations and demonstrates that the frequency of CD107a+ CD4+T cells from DENV immune NHPs, prior to ZIKV infection correlates with enhanced protection from ZIKV challenge. We also noted that DENV specific CD4+T cells isolated one year after DENV infection was highly responsive to the whole DENV virus prior to ZIKV infection (characterized by a significantly higher frequency of IFN-g production and CD107a expression). Also, this group showed a strong trend to higher frequency of reactivity compared to the other two groups, after the other stimulus including the whole ZIKV, ZIKV and DENV envelope and ZIKV nonstructural proteins. Actually, after 30 days of infection the focus of CD4+ T cells reactivity was ZIKV envelope and non-structural antigens. That switch was a trend, but not significant in animals with an early period of convalescence to DENV. Interesting it has been shown that a higher frequency of DENV-specific IFN-g producing T cells are associated with subclinical manifestations in children suffering from secondary DENV infection (60). Our finding on CD4 T cells is consistent with a previous report confirming that preexisting memory CD4 T cells (and not the CD8 T cells or antibodies) are responsible for limiting the severity of illness caused by influenza (61). Notably, the data for CD8+T cells did not recapitulate the observations of the CD4+ T cells. We noted relatively similar responses between the CD8+T cells isolated from the 12M DENV immune compared to the 3M DENV immune animals. This suggests that the CD4+T cell response changed over time leading to potential differences in disease.

The immunopathogenesis of liver damage induced by DENV infection has been addressed in human and animal models, but very limited data is available for ZIKV infection or for sequential DENV/ZIKV or ZIKV/DENV infections. Different immunopathogenic mechanisms like apoptosis, viral replication, autoantibodies to non-structural proteins, infiltrating Natural Killer and CD8+ T cells among others, have been postulated as the intrinsic mechanism of DENV-induced liver damage (62-70). However, there is evidence showing that liver damage is not associated to DENV virus replication *per se* if not with immunopathogenesis induced after the viral infection (63). Fernando et al reported that comparing a well-characterized cohort of 33 cases of non-severe DENV with a group of 22 subjects with severe dengue, the increase of AST, GGT and ALT peaked up by day 6-7 and did not associate with the degree of viremia or the onset or extent of fluid leakage. They found that the liver damage was more related to a possible immune mechanism and associated to higher levels of IL-10 and IL-17 (63). Results from mice also confirm that liver damage is associated with higher systemic levels of proinflammatory cytokines, including TNF-alpha and IL-17, and not to DENV replication (64, 67). In fact, Martinez-Gomez et al showed that anti-TNF-alpha reduce the liver damage significantly without any impact in the viremia (67). Particularly relevant is the limited elevation of the AST in both DENV-immune groups compared to the naïve animals and the increase of that enzyme in 5 of 6 naïve animals 6 days p.i. above the normal ranges. That enzyme (AST) has been reported more frequently elevated and at higher levels than ALT in humans with DENV-induced liver damage (63, 71-74). We found a limited proinflammatory cytokine profile in the naïve animals that may explain the significant increase of liver enzymes in that group. On the other hand, we hypothesize that the limited elevation of liver enzymes in the 3M group at early time point after ZIKV infection is in agreement with the limited immune activation we are reporting in the first days after infection in that group, most likely due to the short period of time between consecutive DENV/ZIKV infections.

The significant role of the T cells in controlling ZIKV replication and as consequence limiting the liver damage in the animals with a DENV-middle convalescence period before ZIKV infection is reinforced by the significant increase of circulating cytolytic protein perforin at day 7 p.i.. We hypothesize that this likely represents T cells acquisition of cytotoxic function (75) in that group compared to the other two groups and correlates with the higher expression of CD107a on the CD4+T cells isolated from the middle convalescent animals. Previously we confirmed a peak in perforin levels in the serum 6 days after ZIKV infection in animals with 2.8 years of previous immunity to DENV (24). Others have shown that Granzyme B levels in CD4+ and CD8+ T cells peaked between 7 and 10 days post-ZIKV infection (50).

The protective role of the cellular immune response controlling the viral burden of ZIKV in mice has been reported (76, 77). More recently mouse models have shown that prior DENV immunity can protect against ZIKV infection during pregnancy, and CD8+ T cells are sufficient for this cross-protection (78). Currently, it is well documented that pre-exposure to DENV both in macaques and humans results in a qualitative modification of the humoral and cellular immune response to ZIKV ^(24, 27-29)^.

The uniqueness of our report is that we provide evidence that the magnitude and the breadth of flavivirus immunity depends not only on pre-infection immune status but time between exposures, with significantly different protection outcomes. Also, taking advantage of the NHP model, we are providing a dissection of the early events of the protective immune response. We are providing mechanistic evidence of the early role of cellular immunity in such protection and characterized a window of transition to the contribution of the humoral immune response.

We are showing that in the presence of previous DENV infection, the increase in the frequency of the specific TEM cells and of the cytotoxic CD4+ T cells and in the magnitude of ZIKV neutralization may occur at any time after ZIKV infection. However, playing a significant role controlling ZIKV viremia and liver damage happens only after a middle-convalescent period of approximately 12 months, but not too early (3 months) or too late (2.8 years) after the primary DENV infection. Interestingly a recent report showed that the IgG3 levels (a marker of recent DENV infection) were positively associated with risk of infection by ZIKV (34). In humans, this fact correlates with our finding in NHPs that a shorter period of DENV immunity may not provide same level of protection against ZIKV replication. Another work reporting results from a children cohort in Nicaragua found that a recent DENV infection was significantly associated with decreased risk of symptomatic ZIKV infection (33). The definition of recent DENV infection was precisely about one year before ZIKV infection which is in agreement with 12M of DENV convalescence we are reporting. We acknowledge that to establish the precise role for the T cells immune response controlling Zika viremia and pathogenesis, depletion of the specific T cells subsets is advised, and our group is already working on that direction.

From our results we cannot anticipate if the effect of previous DENV immunity or the time between DENV and ZIKV infection may have any implications during the pregnancy. More complex studies using a large number of NHPs and well controlled prospective studies in human populations are needed to elucidate such a relationship (79). Based on other results from our group, it is possible to argue that the sequence of ZIKV-DENV infections (41) induce a different immunological response—in terms of the neutralization magnitude, cytokines profile and functionality of the cellular immune response—compared to the DENV-ZIKV scenario shown here. However, in both scenarios, the role of the time interval between infections seems to play a critical role in the quality and quantity of the immune response.

Our findings have enormous impact for the epidemiological models anticipating the magnitude of new ZIKV epidemics in DENV endemic areas and are essential for the planning and evaluation of ZIKV and DENV vaccine schedules, design and monitoring.

## Methods

### Viral stock

ZIKV PRVABC59 strain was obtained from ATCC, BEI Resources (Manassas, VA), was used in order to compare results to our previously published data. This ZIKV strain replicates well in rhesus macaques but has a lower viremia peak than ZIKV H/PF/2013 strain. We aimed to use a strain from the recent epidemic in the Americas region. Virus was expanded and titered by plaque assay and qRT-PCR using protocols standardized in our laboratories. DENV-1 Western Pacific 74, DENV-2 New Guinea 44, DENV-3 Sleman 73 and DENV-4 Dominique strains kindly provided by Steve Whitehead (National Institutes of Health, Bethesda, Maryland) were used for neutralization assays. DENV-2 New Guinea 44 strain was also used to infect macaques in September 2016 and June 2017.

### Ethics Statement

All procedures were reviewed and approved by the Institute’s Animal Care and Use Committee at Medical Sciences Campus, University of Puerto Rico (IACUC-UPR-MSC) and performed in a facility accredited by the Association for Assessment and Accreditation of Laboratory Animal Care (AAALAC) (Animal Welfare Assurance number A3421; protocol number, 7890116). Procedures involving all study animals were approved by the Medical Sciences Campus, UPR IACUC and were conducted in accordance with USDA Animal Welfare Regulations, the Guide for the Care and use of Laboratory Animals and institutional policies. In addition, steps were taken to lighten sufferings, including use of anesthesia and method of sacrifice if appropriate, in accordance with the recommendations of the Guide for the Care and use of Laboratory Animals (8th edition), Animal Welfare Act and the Public Health Service (PHS) Policy on Humane Care and Use of Laboratory Animals and in accordance with the recommendations of the Weatherall report, “The use of non-human primates in research: http://www.acmedsci.ac.uk/more/news/the-use-of-non-human-primates-in-research/. Macaques were continuously monitored by trained veterinarians at the Animal Research Center and evaluated twice daily for evidence of disease or injury. Feeding and drinking continued normally during this period. All procedures were conducted under anesthesia by intramuscular injection of ketamine at 10–20 mg/kg-1 of body weight, as approved by the IACUC. Anesthesia was delivered in the caudal thigh using a 23-Gauge sterile syringe needle. During the period of the entire study, the macaques were under the environmental-enrichment program of the facility, also approved by the IACUC.

### Immunization and virus challenge of macaques

Young adult rhesus macaques (4-7 years of age) seronegative for DENV and ZIKV were housed in the CPRC facilities, University of Puerto Rico, San Juan, Puerto Rico. For the ZIKV challenge, macaques previously infected with DENV-2 in September 2016 (Cohort 1, n=6) and June 2017 (Cohort 2, n=4), and DENV/ZIKV-naïve macaques (Cohort 3, n=6) were infected subcutaneously in the deltoid area with 500uL of virus diluted in PBS, using a dose of 1 × 10^6^ pfu. All macaques were male. The average age for cohort 1 was 5.1 years (5.6, 4.9, 5.1, 4.9, 5.0 and 5.0 years), 6.8 years for cohort 2 (5.5, 7.75, 7.6 and 6.7 years), and 5.7 years for cohort 3 (6.3, 6.5, 5.8, 4.75, 5.4 and 5.6 years). Weights were taken on Day 0 and every other day during the acute infection period (days 1–6, then at day 30 p.i.). Rectal and external temperature were taken daily during the acute infection period (days 1–6 and on day 30 p.i.). External temperature was recorded using an infrared device (EXTECH Instruments, Waltham, MA) as per the manufacturer’s instructions. Blood and chemical tests were performed on day 0 and every other day until day 6 p.i.

### Note on sample collection

The intended schedule was unexpectedly affected by Hurricane María. Sample collection programmed from days 7 to 29 p.i. was interrupted due to inability of access and/or lack of electricity in CPRC facilities in University of Puerto Rico, San Juan, Puerto Rico.

### DENV and ZIKV titration and neutralization assays

For virus titration, Vero81 cells (ATCC CCL-81) at approx. 8.5 × 10^4^ cells/well in 24 well plates with growth medium (Dulbecco’s Modified Eagle’s medium, Thermo Fisher Scientific, with 10% FBS (Gibco), 1% non-essential amino acids (Gibco), 1% HEPES (Gibco), 1% L-glutamine (Gibco) and 1% Pen/Strep (Gibco)). At around 85% confluency (approx. 24 hours later), ten-fold dilutions of virus were prepared in diluent medium (Opti-MEM (Invitrogen) with 2% FBS (Gibco) and 1% antibiotic/antimycotic (HyClone) and added to the wells after removing growth medium. Each virus dilution was added in 100 mL triplicates, and plates were incubated 1 hour at 37C/5%CO2/rocking. After incubation, 1 mL of overlay (Opti-MEM with 1% Carboxymethylcellulose (Sigma), 2% FBS, 1% non-essential amino acids (Gibco), 1% antibiotic/antimycotic (HyClone)) was added to the plates containing viral dilutions, followed by an incubation period at 37C/5%CO2. After 3 to 5 days of incubation (depending of the virus), overlay was washed away with phosphate buffered saline (PBS) and fixed with 80% methanol. For ZIKV, cells were stained with crystal violet after fixing. For DENV, plates were fixed then blocked with 5% non-fat dry milk in PBS and incubated for 1hr/37C/5%CO2/rocking with anti-E mAb 4G2 and anti-prM mAb 2H2 (provided by Dr. Aravinda de Silva), both diluted 1:250 in blocking buffer. Plated were washed twice and incubated 1hr/37C/5%CO2/rocking with horseradish peroxidase (HRP)-conjugated goat anti-mouse antibody (Sigma), diluted 1:1,000 in blocking buffer. Foci were developed with TrueBlue HRP substrate (KPL) and counted. For the Focus/Plaque Reduction Neutralization Test (FRNT/PRNT), sera were diluted two-fold and mixed with approx. 35 foci per plaque-forming units (FFU per p.f.u. per mL) of virus and then incubated for 1hr/37C/5%CO2/rocking. Virus-serum dilutions were added to 24 well-plates containing Vero81 cells as mentioned above, and incubation was continued for 1hr/37C/5%CO2/rocking. After incubation, overlay was added, and the aforementioned procedure was repeated. Mean focus diameter was calculated from approx. 20 foci per clone measured at X5 magnification. Results were reported as the FRNT or PRNT with 60% or greater reduction in DENV or ZIKV foci or plaques (FRNT60 or PRNT60). A positive neutralization titer was designated as 1:20 or greater, while <1:20 was considered a negative neutralization titer.

### qRT-PCR

Viral RNA for real-time PCR assay was extracted from 140 ml of virus isolate (previously tittered as described above) and serum samples using Invitrogen PureLink RNA Mini Kit (Invitrogen, Valencia, CA) as per the manufacturer’s instructions. Real-time RT-PCR (TaqMan) assay-specific primers and probes for ZIKV were designed by Sigma-Aldrich (St Louis, MO) following the protocol developed by the Molecular Diagnostics and Research Laboratory Centers for Disease Control and Prevention (CDC), Dengue Branch at San Juan, PR. RNA from other flaviviruses were included as negative control. For the reaction mixture, 5 ml of RNA was combined with 100 mM primers and 25 mM probe in a 25 ml total volume using Life Technologies SuperScriptIII Platinum assay kit (Life Sciences). Assays were performed in an iCycler iQ Real Time Detection System (Bio Rad, CA). For quantification, a standard curve was generated from ten-fold dilutions of RNA from a known amount of virus.

### ELISA for DENV and ZIKV

Prior to ZIKV challenge, DENV/ZIKV seronegative status of cohort 3 animals was assessed using DENV IgG/IgM and ZIKV NS1 IgG commercial kits (Focus Diagnostics, CA). After ZIKV infection, seroreactivity to DENV was tested using commercial IgG and IgM ELISA kits (Focus Diagnostics, Cypress, CA). ZIKV IgG was assessed with available commercial kits (XpressBio, Frederick, MD and InBios, Seattle, WA respectively). ZIKV-NS1 IgG was examined using a commercial kit (Alpha Diagnostic, San Antonio, TX). All tests were performed per the manufacturer’s instructions. For the measurement of IgM levels against ZIKV, samples were tested using a ZIKV IgM MAC-ELISA assay developed by Aravinda de Silva’s laboratory. Briefly, a 96-well microtiter plate was coated with anti-human IgM (1:50). The plate was left at 4 C° overnight. Following incubation, coating was removed by dumping and the plate was blocked for 30 minutes at room temperature. After blocking, the plate was washed once, and sample dilutions were prepared (1:40) and added to the plate. Positive and negative controls for ZIKV and DENV were also prepared. After the addition of samples and controls, the plate was incubated one hour at 37 C° in a humidified chamber. Before adding the antigens, the plate was washed twice. Stock C6/36 ZIKV and DENV antigens were diluted (1:2 and 1:3, respectively), and added to the plate. The plate was incubated overnight at 4 C° in a humidified chamber. The next day, the plate was washed twice, and a horseradish peroxidase (HRP)-conjugated monoclonal antibody (6B6C-1) was added, followed by incubation for one hour at 37 C° in a humidified chamber. Detecting antibody was diluted in blocking buffer 1:1000 prior to addition to plate. A last cycle of washing (twice) was performed, and TMB substrate was added to all wells. Plate was covered immediately to block out light and incubated at room temperature for 30 minutes. Before colorimetric detection, the plate was allowed to sit at room temperature for 5 minutes. Optical density at 450 nm (OD) values were measured in three separate readings at 5-minute intervals. Results were expressed as mean OD of sample reacted with viral antigen (P)/mean OD of normal human serum reacted with viral antigen (N) and reported as negative (P/N value of <2), presumptive positive (P/N value of >3) or equivocal (2< P/N <3).

### Immunophenotyping

Phenotypic characterization of rhesus macaque PBMCs was performed by multicolor flow cytometry using direct immunofluorescence. Aliquots of 150 ul of heparinized whole blood were directly incubated with a mix of antibodies for 30 min. at room temperature. After incubation, red blood cells were fixed and lysed with ACK, and cells were washed three times with PBS. Samples were analyzed using a MACSQuant Analyzer 10 flow cytometer (Miltenyi Biotec, CA). The following antibodies were used in this study: CD123-APC (7G3), CD20-FITC (2H7), CD14-FITC (M5E2), CD16-Alexa Fluor 700 (3G8), CD20-PacBlue (2H7), CD69-PE (FN50), CD14-V500 (M5E2), KI67-FITC (B56) and CD3-FITC (SP34) from BD-Biosciences; CD4-PerCP (M-T466), HLA-DR VioGreen (G46.6), CD337 (NKp30)-PE-Vio770 (AF29-4D12), CD8-VioGreen (BW135/80), CD159a (NKG2A)-FITC (REA110), CD3-PE-Vio770 (10D12), CD16-APC-Vio770 (VEP13), CD3-APC (10D12), CD28-APC-Vio770 (15E8) and CD56-PE (AF12-7H3) from Miltenyi; CD335 (NKp46)-PC5 (BAB281) from Beckman-Coulter; CD11c-PE/Cy7 (3.9), CD8-FITC (SK1) and CD8-BV421 (SK1) from Biolegend. For analyses, LYM were gated based on their characteristic forward and side scatter pattern; T cells were then selected with a second gate on the CD3 positive population. CD8+ T cells were defined as CD8^+^CD3^+^ and CD4+ T cells were CD4^+^CD3^+^. Natural Killer cells were defined as CD3^-^CD20^-^ CD14^-^ and analyzed by the expression of NK cell markers CD16, CD8, NKG2A, NKG2C, NKp30 and NKp46. B cells were defined as CD20^+^CD3^-^CD14^-^. Activation marker CD69 was determined in each different lymphoid cell population. Monocytes were defined as CD20^-^CD3^-^CD14^+^ and CD20^-^CD3^-^CD14^+^CD16^+^. Finally, dendritic cells (DCs) were separated into two populations by the expression of CD123 (pDCs) or CD11c (mDCs) in the HLA^-^DR^+^CD3^-^ CD14^-^CD20^-^ population. Data analysis was performed using Flowjo (Treesar).

### Cellular immune response analysis

Intracellular cytokine staining of PBMCs from rhesus macaques was performed by multicolor flow cytometry using methods similar to those described by Meyer et al. Briefly, PBMC samples were thawed 1 day prior to stimulation. Approx. 1.5 × 10^6^ PBMCs were infected overnight with DENV-2 (NGC44) at a MOI of 0.1 or ZIKV at a MOI of 0.5 in RPMI medium with 5% FBS. The remaining PBMCs were rested overnight as described earlier in 5ml of RPMI with 10% FBS. These PBMCs were then stimulated for 6 h at 37C/5%/CO2 with ZIKV-E peptides (15-mers overlapping by 10 amino acids, 2.5 ug/ml^-1^ per peptide), ZIKV-NS1 protein peptides (15-mers overlapping by 10 amino acids, 475 ng/ml^-1^ per peptide), or DENV-2 E peptides (1.25 ug/ml^-1^), all in the presence of brefeldin A (10 ug/ml^-1^), a-CD107a-FITC (H4A3) (10 ul), and co-stimulated with a-CD28.2 (1 ug/ml^-1^) and a-CD49d (1 ug/ml^-1^). After stimulation, the cells were stained for the following markers: CD4-PerCP Cy5.5 (Leu-3A (SK3), CD8b-Texas Red (2ST8.5H7), CD3-PacBlue (SP34), CD20-BV605 (2H7), CD95-V510 (DX2), CD28.2-PE-Cy5, IFN-g-APC (B27) and TNF-a-PE-Cy7 (MAB11). The samples were run on an LSRII (BD) and analyzed using Flowjo (Treesar). Lymphocytes were gated based on their characteristic forward and side scatter pattern, T cells were selected with a second gate on the CD3-positive population, and at the same time CD20 positive cells were excluded. CD8+ T cells were defined as CD3^+^ CD20^-^CD8^+^ and CD4+ T cells as CD3^+^CD20^-^ CD4^+^. Cytokine expression was determined by the per cent CD4+ or CD8+ positive cells, and then stained positive for the cytokine IFN-g or TNF-a. CD107a were also measured in these populations to determine functional cytotoxicity. Further analysis was also performed to examine CD28 and CD95 expression on the LYM populations to study the presence of central and effector memory cell populations.

### Multiplex cytokine analysis

Sera from rhesus macaques was analyzed for 14 cytokines and chemokines by Luminex using established protocols for Old World primates. Evaluation of analytes B cell-activating factor (BAFF), eotaxin (CCL11), interferon alpha (IFN-a), IFN-g, IL-1 receptor antagonist (IL-1Ra), IL-6, interferon-inducible T-cell alpha chemoattractant (I-TAC, CXCL11), monocyte chemoattractant protein 1 (MCP-1, CCL2), macrophage migration inhibitory factor (MIF), monokine induced by gamma interferon (MIG, CXCL9), inflammatory protein 1-alpha (MIP-1a, CCL3), MIP-1b (CCL4), perforin and interferon gamma-induced protein 10 (IP-10, CXCL10) were included in this assay.

### Statistical methods

Statistical analyses were performed using GraphPad Prism 7.0 software (GraphPad Software, San Diego, CA, USA). For viral burden analysis, the log titers and levels of vRNA were analyzed unpaired multiple t tests and two-way ANOVA. Also, a Chi-squared test was used to analyze a contingency table created from obtained viremia data. The statistical significance between or within groups evaluated at different time points was determined using two-way analysis of variance (ANOVA) (Tukey’s, Sidak’s or Dunnett’s multiple comparisons test) or unpaired t-test to compare the means. The p values are expressed in relational terms with the alpha values. The significance threshold for all analyses was set at 0.05; p values less than 0.01 are expressed as P<0.01, while p values less than 0.001 are expressed as P<0.001. Similarly, values less than 0.005 are expressed as P<0.005. In figures, p values from 0.01 to 0.05 are depicted as *, 0.001 to 0.01 as **, 0.0001 to 0.001 as ***, and lastly, values less than 0.0001 are depicted as ****.

## Data availability

All relevant data are available from the authors upon request.

## Acknowledgements

This work would have not been possible without the dedication and commitment of the Caribbean Primate Research Center and the Animal Resources Center staff. We also acknowledge the efforts of all persons that in the aftermath of the hurricane Maria showed a high level of resilience making possible to advance the science in awkward situations. Authors recognize the support provided by Dr. Elmer Rodriguez reviewing the statistics and by Dr. Willy Ramos for the useful review of the immunology data. This work received support from grant by Grants 2 P40 OD012217 and 2U42OD021458-15 to MIM and CAS (ORIP, OD, NIH), and R25GM061838 to C.S.-C. Also, partial support was provided by Grant K22AI104794 to JDB (NIAID) and partially used resources that were supported by the Southwest National Primate Research Center grant P51 OD011133 from the Office of Research Infrastructure Programs, National Institutes of Health.

