## Supplementary figures for "Significant control of Zika infection in macaques depends on the elapsing time after dengue exposure"

**Supporting Information**

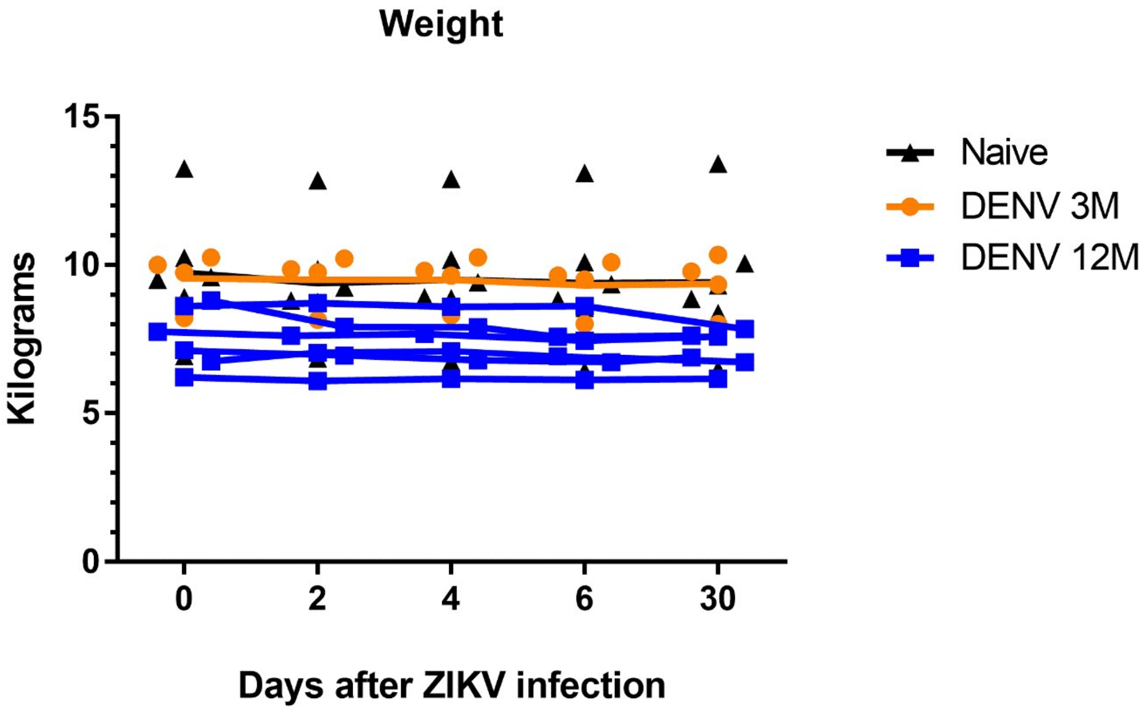


**S1 Figure. Weight distribution per animal cohort.** Weight of rhesus macaques is expressed in kilograms (kg). Animals exposed to DENV 12 months before ZIKV infection are depicted in blue, while animals exposed to DENV 3 months before are in orange. Naïve animals are in black.

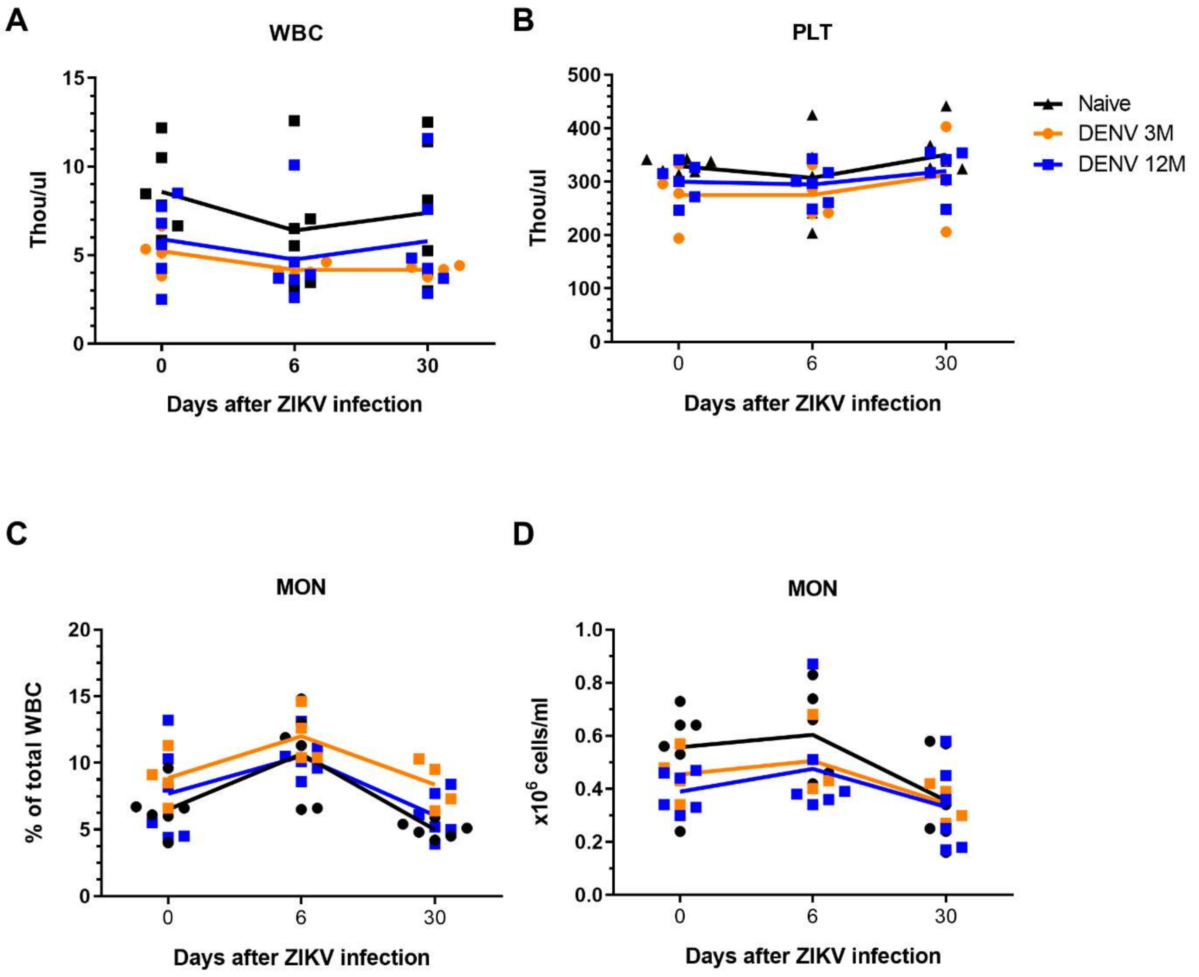


**S2 Figure. Kinetics of hematology and laboratory results.** Cell subsets obtained from complete blood count (CBC) tests at baseline, 6 and 30 days p.i. In all panels, animals exposed to DENV 12 months before ZIKV infection are in blue, while animals exposed to DENV 3 months before are in orange. Naïve animals are in black. (A) White blood cells (WBC) total depicted in thou/uL. (B) Platelet (PLT) levels total depicted in thou/uL. (C-D) Monocyte (MON) kinetics expressed as percentage of total WBC and absolute numbers (x10^6 cells/mL).

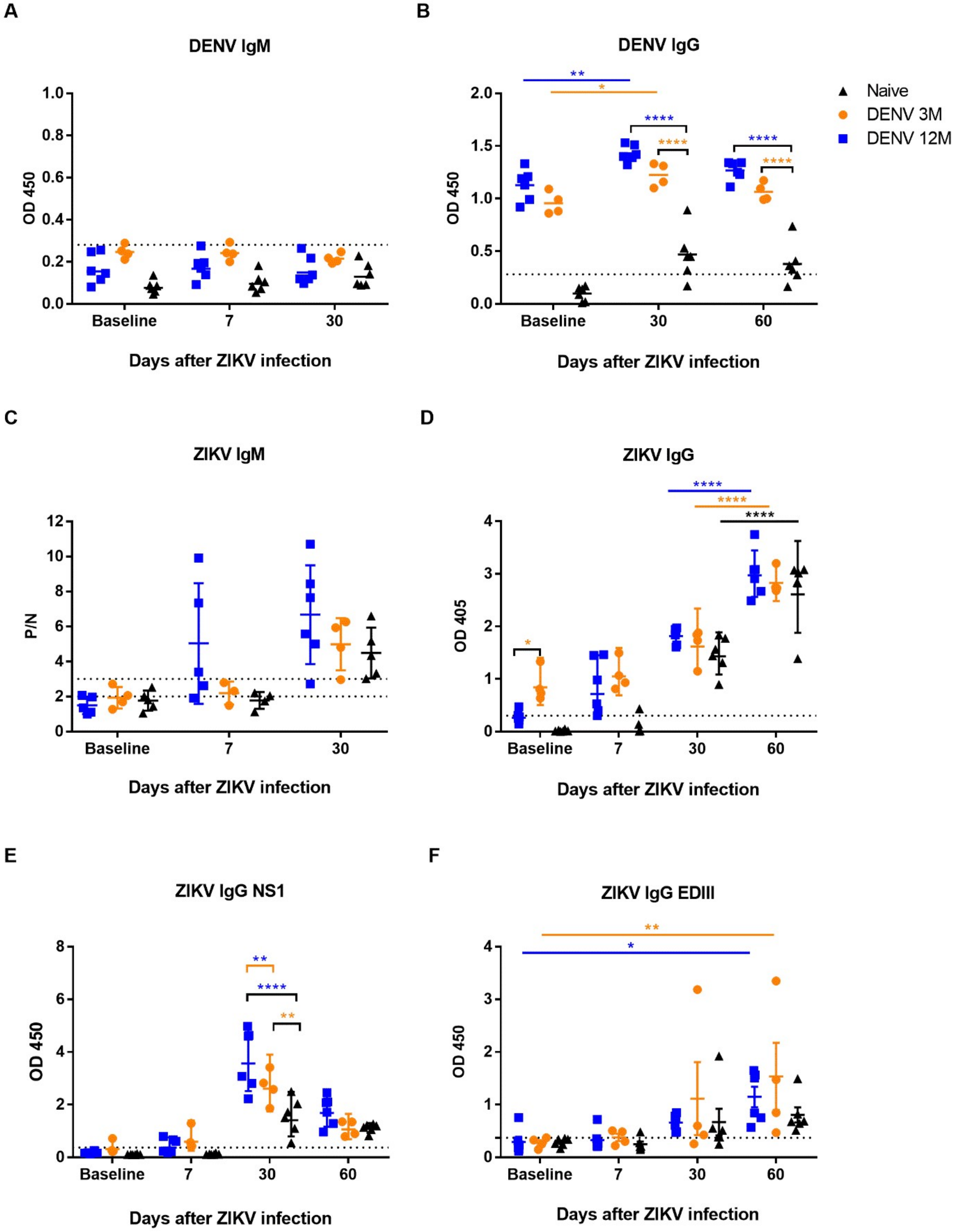

A
DENV IgM
OD 450
1.0
0.8
0.6
0.4
0.2
0.0
Baseline
7
30
Days after ZIKV infection
B
DENV IgG
OD 450
2.0
1.5
1.0
0.5
0.0
Baseline
30
60
Days after ZIKV infection
Naive
DENV 3M
DENV 12M
C
ZIKV IgM
P/N
12
10
8
6
4
2
0
Baseline
7
30
Days after ZIKV infection
D
ZIKV IgG
OD 405
4
3
2
1
0
Baseline
7
30
60
Days after ZIKV infection
E
ZIKV IgG NS1
OD 450
8
6
4
2
0
Baseline
7
30
60
Days after ZIKV infection
F
ZIKV IgG EDIII
OD 450
4
3
2
1
0
Baseline
7
30
60
Days after ZIKV infection

**S3 Figure. Serological profile of the three cohorts of macaques before and after ZIKV infection.** Humoral response was assessed using different commercial and in-house ELISA tests. (A-F) Binding capacity of antibodies from animals with different immune background are shown. Animals from cohort 1 are shown in blue, animals from cohort 2 are shown in orange and naïve animals from cohort 3 are shown in black in all panels. Dotted lines indicate the limit of detection for each test. For panel C, samples below lowest dotted line are negative (PN value <2), samples above the highest dotted line are presumptive positive (P/N value >3), and samples between the two dotted lines are equivocal (2< P/N <3). Statistically significant differences among and within groups were calculated by two-way ANOVA using Tukey's multiple comparisons test (*P<0.05, **P<0.005 and ****P<0.0001). Colored stars represent a significantly different group, while colored lines represent the group that it is compared to.

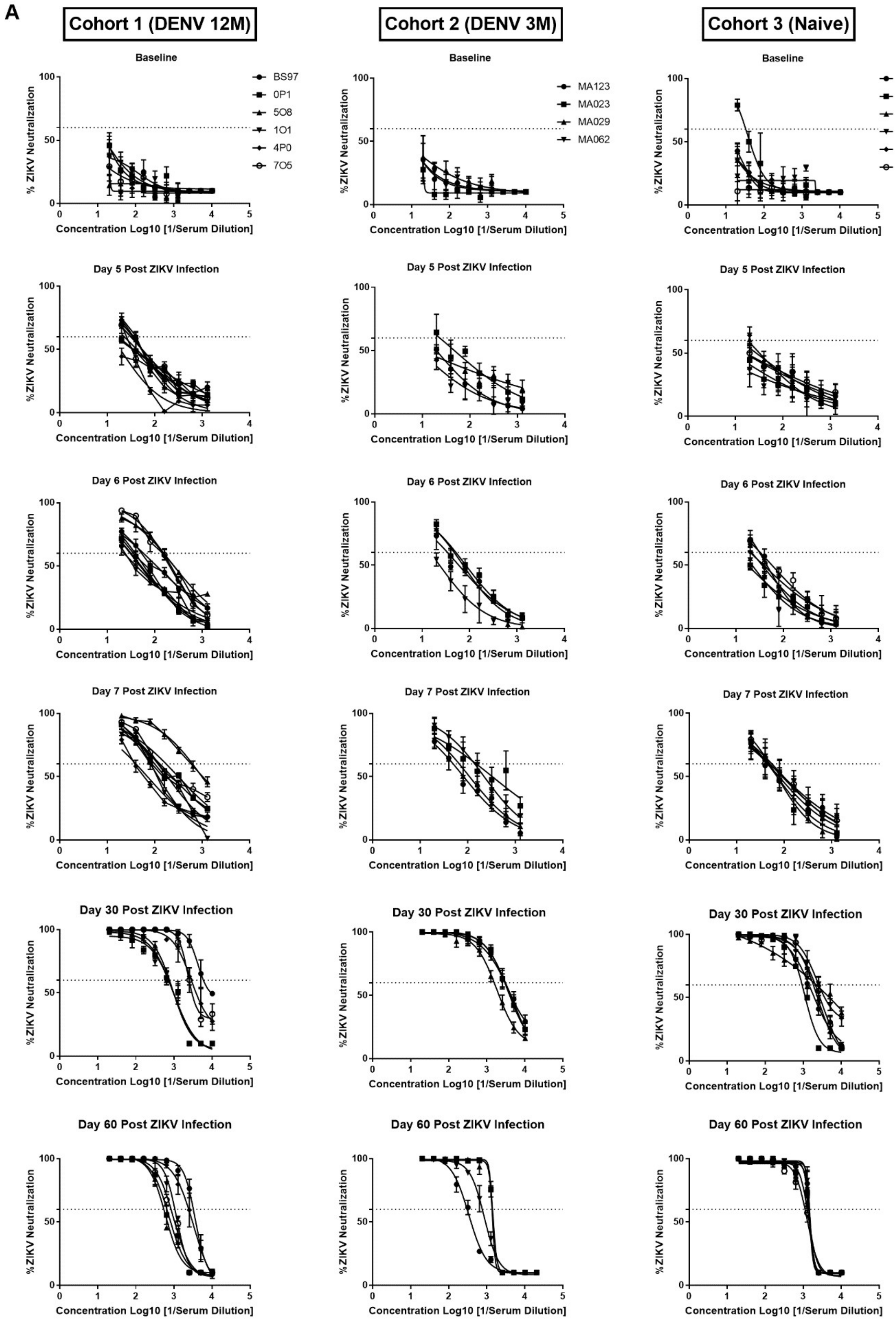

A
Cohort 1 (DENV 12M)
Cohort 2 (DENV 3M)
Cohort 3 (Naive)
Baseline
BS97
0P1
5O8
1O1
4P0
7O5
MA123
MA023
MA029
MA062
MA067
MA068
BZ34
MA141
MA143
MA085
Day 5 Post ZIKV Infection
Day 6 Post ZIKV Infection
Day 7 Post ZIKV Infection
Day 30 Post ZIKV Infection
Day 60 Post ZIKV Infection
% ZIKV Neutralization
Concentration Log10 [1/Serum Dilution]

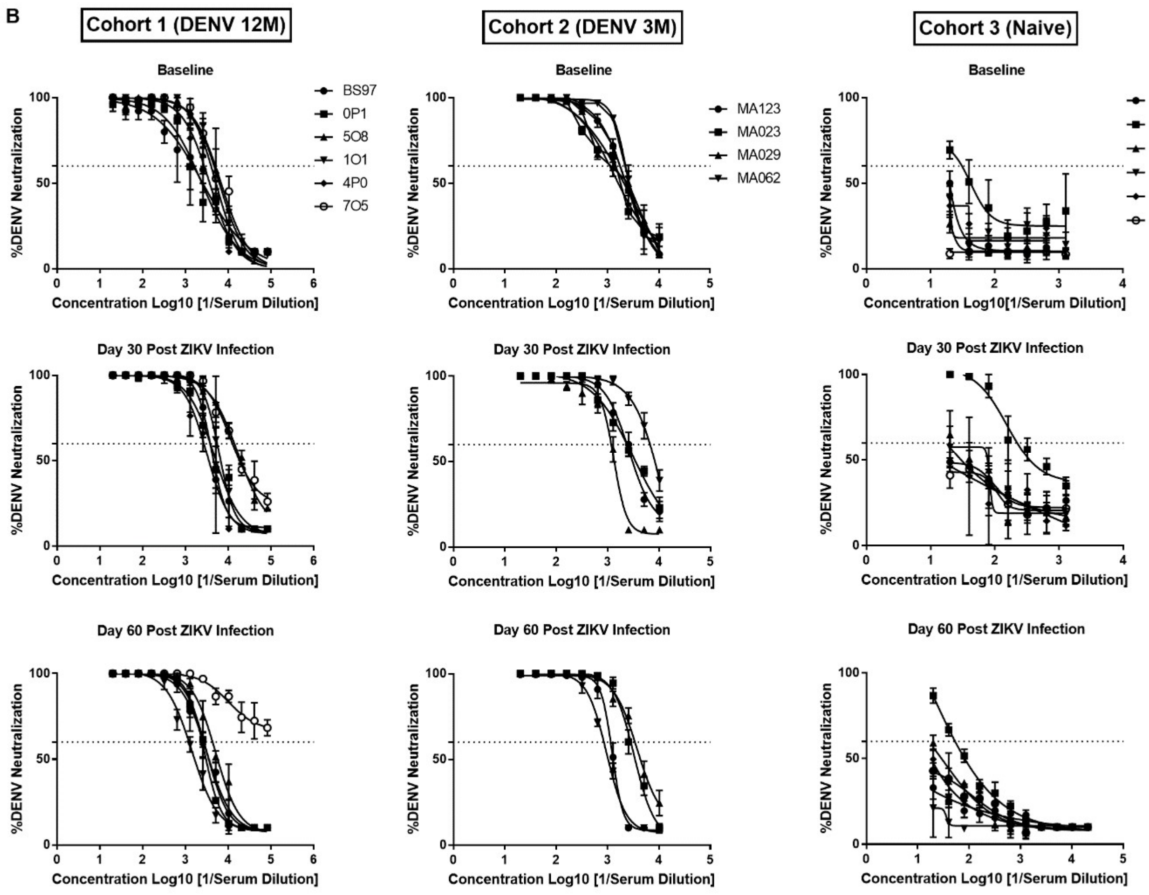


**S4 Figure. Dynamic of the neutralizing antibodies to ZIKV and DENV.** The relationship between dilution titers and the magnitude of neutralizing antibodies before and after ZIKV infection is shown. (A) Neutralization magnitude against ZIKV before, and at days 3, 5, 6, 7, 30 and 60 after ZIKV infection was determined using a dilution:neutralization capacity relation. (B) Neutralization magnitude against DENV 2 before, and 30 and 60 days after ZIKV infection was determined using a dilution:neutralization capacity relation.

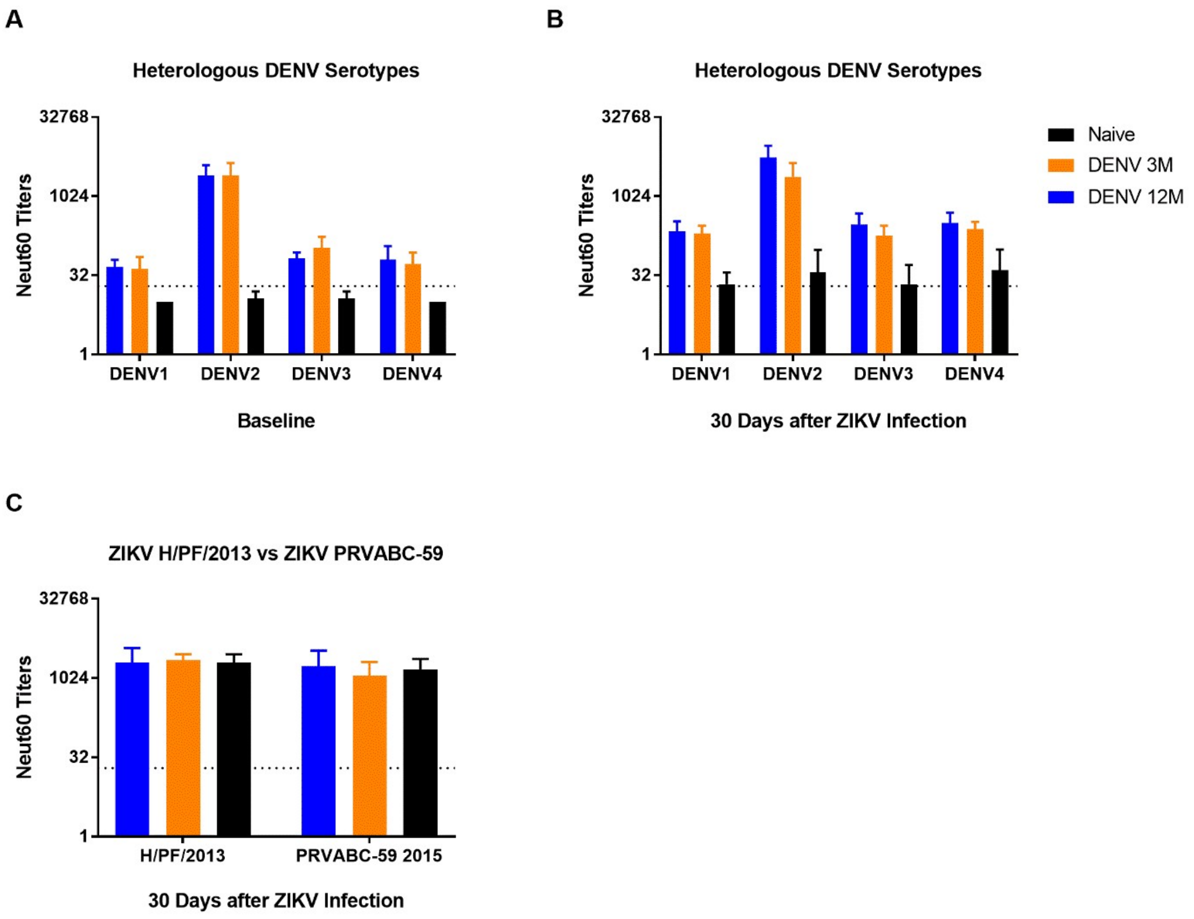


**S5 Figure. Neutralizing response to heterologous DENV serotypes and two different ZIKV strains.** PRNT and FRNT assays were performed to determine the effect of previous DENV immunity in a subsequent ZIKV infection, and the neutralizing antibody response against different dengue serotypes and zika strains. In all panels, animals exposed to DENV 12 months before ZIKV infection are in blue, while animals exposed to DENV 3 months before are in orange. Naïve animals are in black. (A-B) Neutralizing response against heterologous DENV serotypes before and after ZIKV infection. (C) Neutralization against two different ZIKV strains was performed. Dotted lines indicate the limit of detection for the assay.

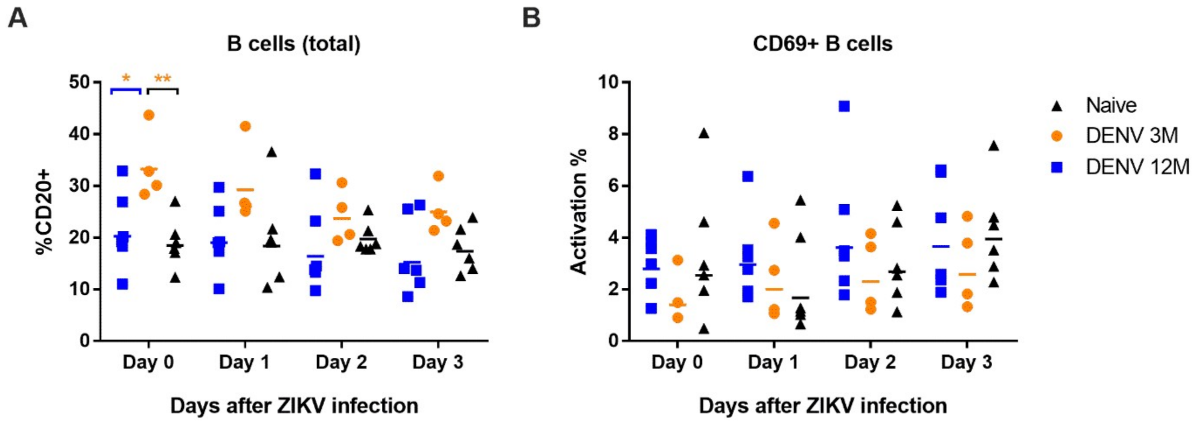


**S6 Figure. B cells profile before and after ZIKV infection.** Frequency of B cells was assessed. In all panels, animals exposed to DENV 12 months before ZIKV infection are in blue, while animals exposed to DENV 3 months before are in orange. Naïve animals are in black. (A) Percentage of total B cells (CD20+) during baseline and days 1 through 3 p.i. (B) Frequency of activated B cells (CD20+ CD69+) during baseline and days 1 through 3 p.i. Comparisons between cohorts were performed by two-way ANOVA using Tukey's multiple comparisons test (*$P<0.05$ and **$P<0.01$). Colored stars represent a significantly different group, while colored lines represent the group that it is compared to.

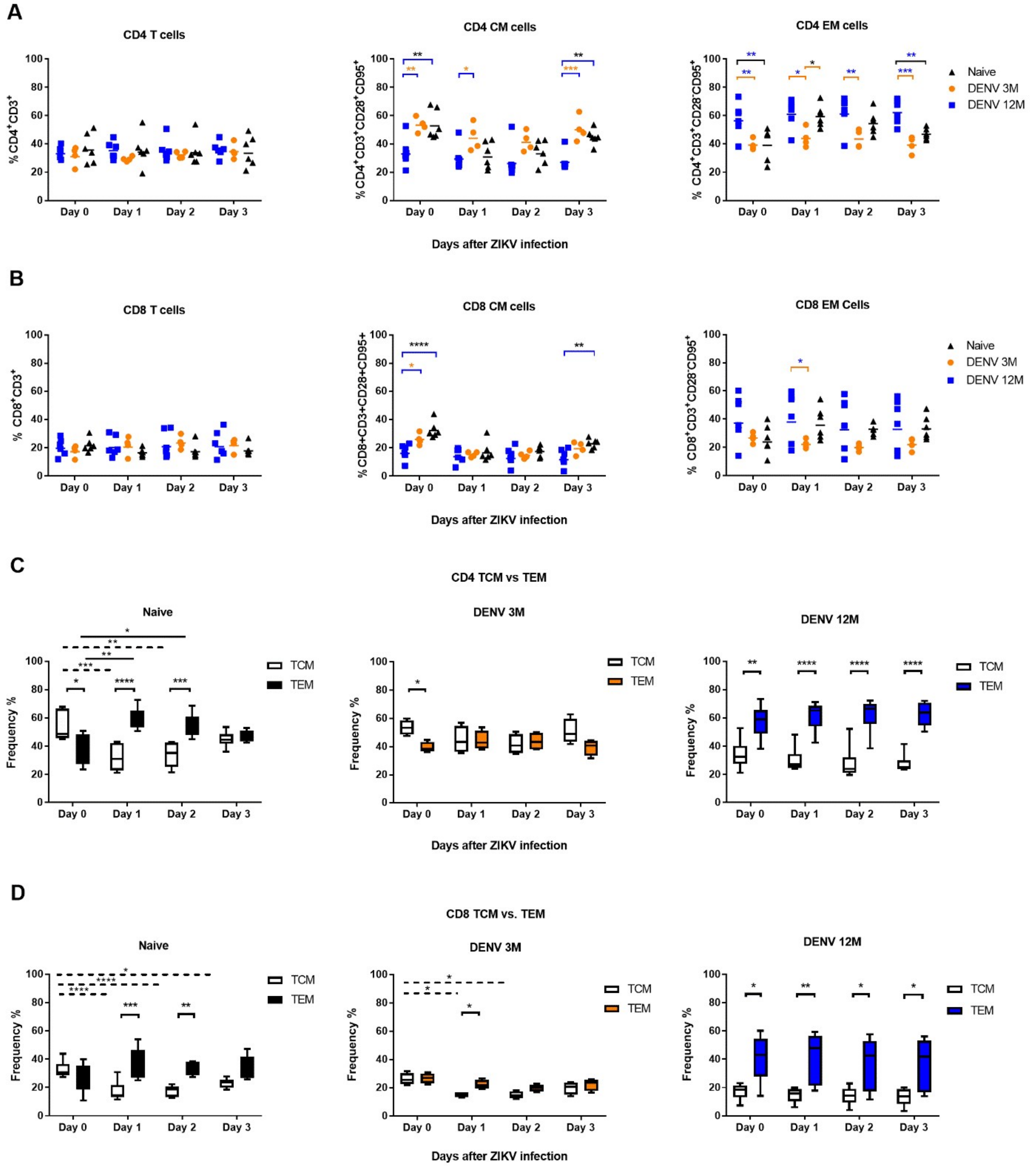


**S7 Figure. Dynamic and relationship of central and effector memory cells in the CD4⁺ and CD8⁺ T compartments.** Changes in the level of and in the relationship between the central and effector memory cells are modified by the time passed after a previous DENV immunity. In all panels, animals exposed to DENV 12 months before ZIKV infection are in blue, while animals exposed to DENV 3 months before are in orange. Naïve animals are in black. (A) Frequency of CD4 T cells, central memory CD4 T cells and effector memory CD4 T cells. (B) Frequency of CD8 T cells, central memory CD8 T cells and effector memory CD8 T cells. (C) Frequency of CD4

TCM and TEM cells in each group separately. (D) Frequency of CD8 TCM and TEM cells in each group separately. Dotted and solid lines denote statistical differences between TCM or TEM cells at different time points respectively. Statistically significant differences among groups were calculated by two-way ANOVA using Tukey's multiple comparisons test (*$P<0.05$). Colored stars represent a significantly different group, while colored lines represent the group that it is compared to.

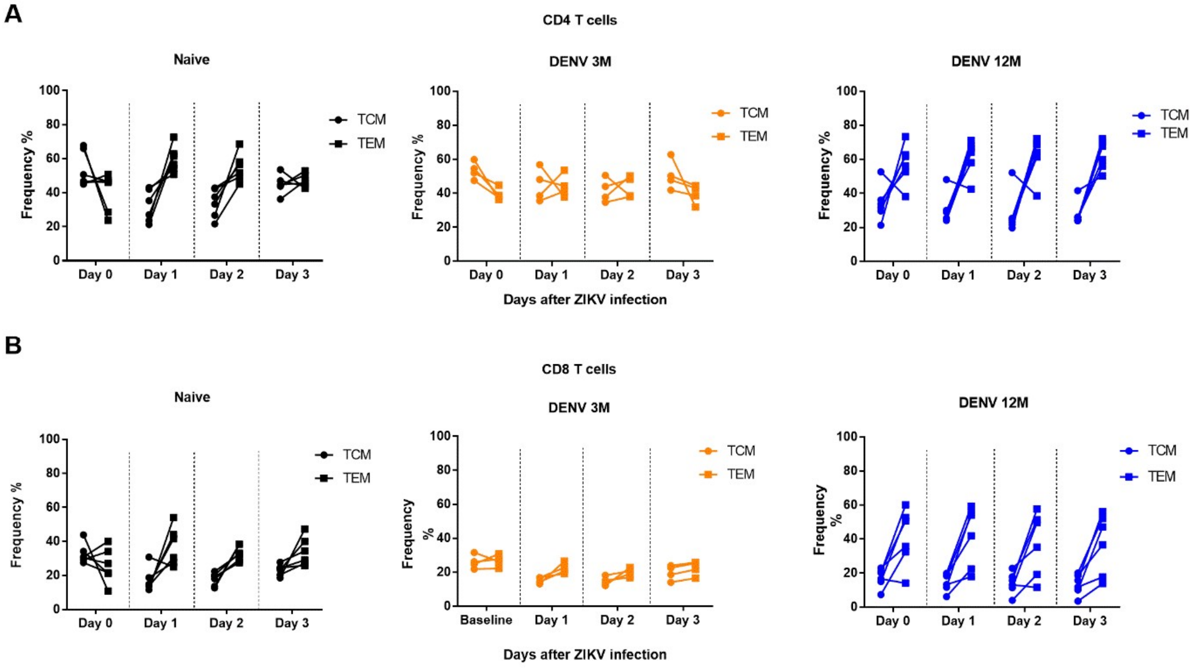


**S8 Figure. Changes in frequency of central and effector memory cells per cohort after ZIKV infection.** Displacement from a central to an effector subset in CD4 and CD8 T cells is shown. Naïve animals are depicted in black. Animals exposed to DENV 12 months before ZIKV infection are in blue, while animals exposed to DENV 3 months before are in orange. (A) Frequency of CD4 TCM and TEM cells in days 0 (before infection) through 3 p.i. (B) Frequency of CD8 TCM and TEM cells in days 0 (before infection) through 3 p.i.

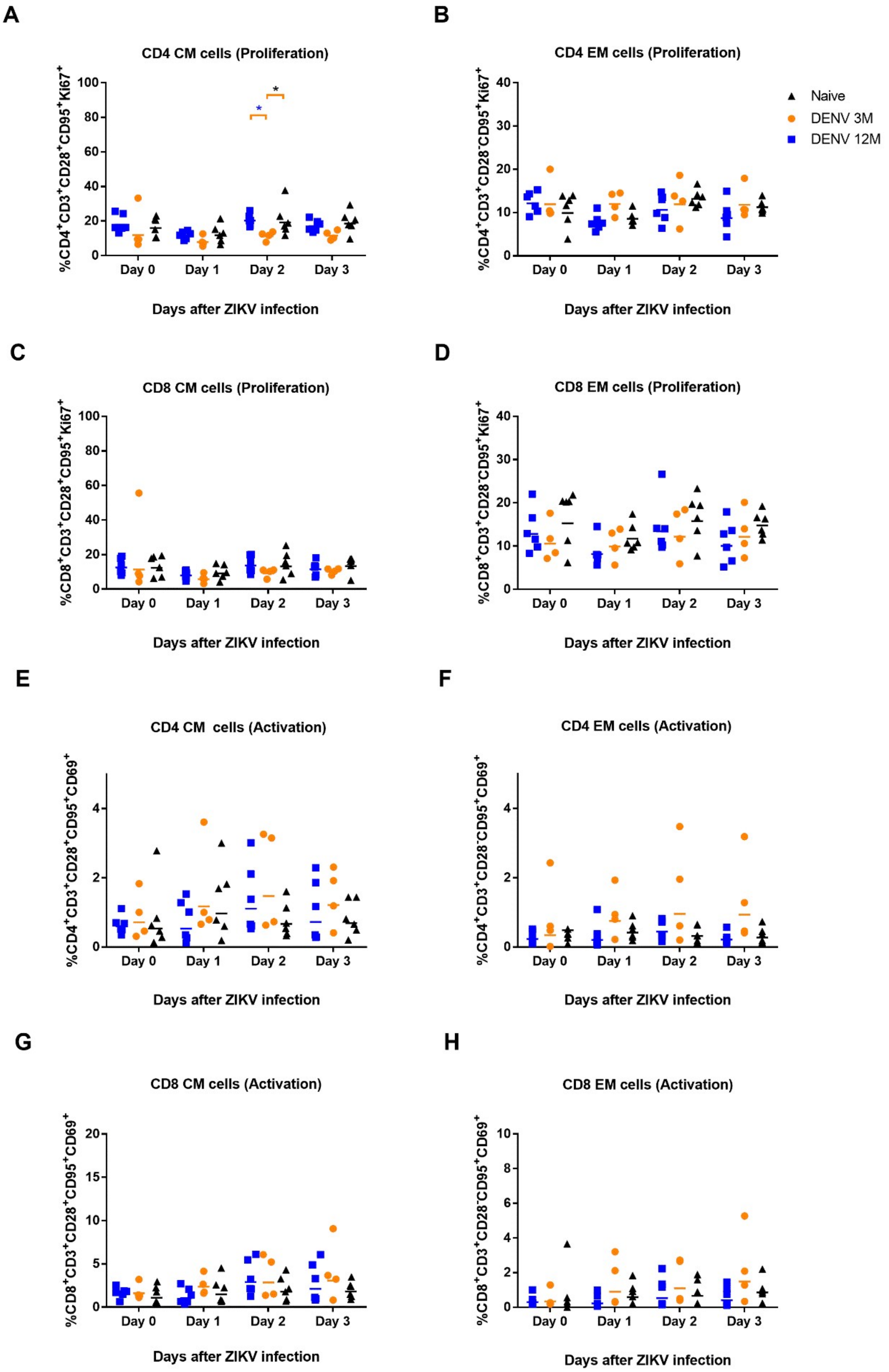
A
CD4 CM cells (Proliferation)
%CD4+CD3+CD28+CD95+Ki67+
Days after ZIKV infection
B
CD4 EM cells (Proliferation)
%CD4+CD3+CD28-CD95+Ki67+
Days after ZIKV infection
Naive
DENV 3M
DENV 12M
C
CD8 CM cells (Proliferation)
%CD8+CD3+CD28+CD95+Ki67+
Days after ZIKV infection
D
CD8 EM cells (Proliferation)
%CD8+CD3+CD28-CD95+Ki67+
Days after ZIKV infection
E
CD4 CM cells (Activation)
%CD4+CD3+CD28+CD95+CD69+
Days after ZIKV infection
F
CD4 EM cells (Activation)
%CD4+CD3+CD28-CD95+CD69+
Days after ZIKV infection
G
CD8 CM cells (Activation)
%CD8+CD3+CD28+CD95+CD69+
Days after ZIKV infection
H
CD8 EM cells (Activation)
%CD8+CD3+CD28-CD95+CD69+
Days after ZIKV infection
Day 0
Day 1
Day 2
Day 3

**S9 Figure. Proliferation and activation profile of central and effector memories cells related to time of ZIKV infection.** CD4 and CD8 T cell central and effector memory subsets were evaluated in terms of proliferation and activation. Animals from cohort 1 are depicted in blue, while animals from cohort 2 are shown in orange. Naïve animals from cohort 3 are shown in black. (A-B) Proliferation levels of CD4 TCM and TEM cells before (Day 0) and after (Days 1 through 3) ZIKV infection. (C-D) Proliferation levels of CD8 TCM and TEM cells before (Day 0) and after (Days 1 through 3) ZIKV infection. (E-F) Activation levels of CD4 TCM and TEM cells before (Day 0) and after (Days 1 through 3) ZIKV infection. (G-H) Activation levels of CD8 TCM and TEM cells before (Day 0) and after (Days 1 through 3) ZIKV infection. Comparisons between cohorts were performed by two-way ANOVA using Tukey's multiple comparisons test (*P<0.05). Colored stars represent a significantly different group, while colored lines represent the group that it is compared to.

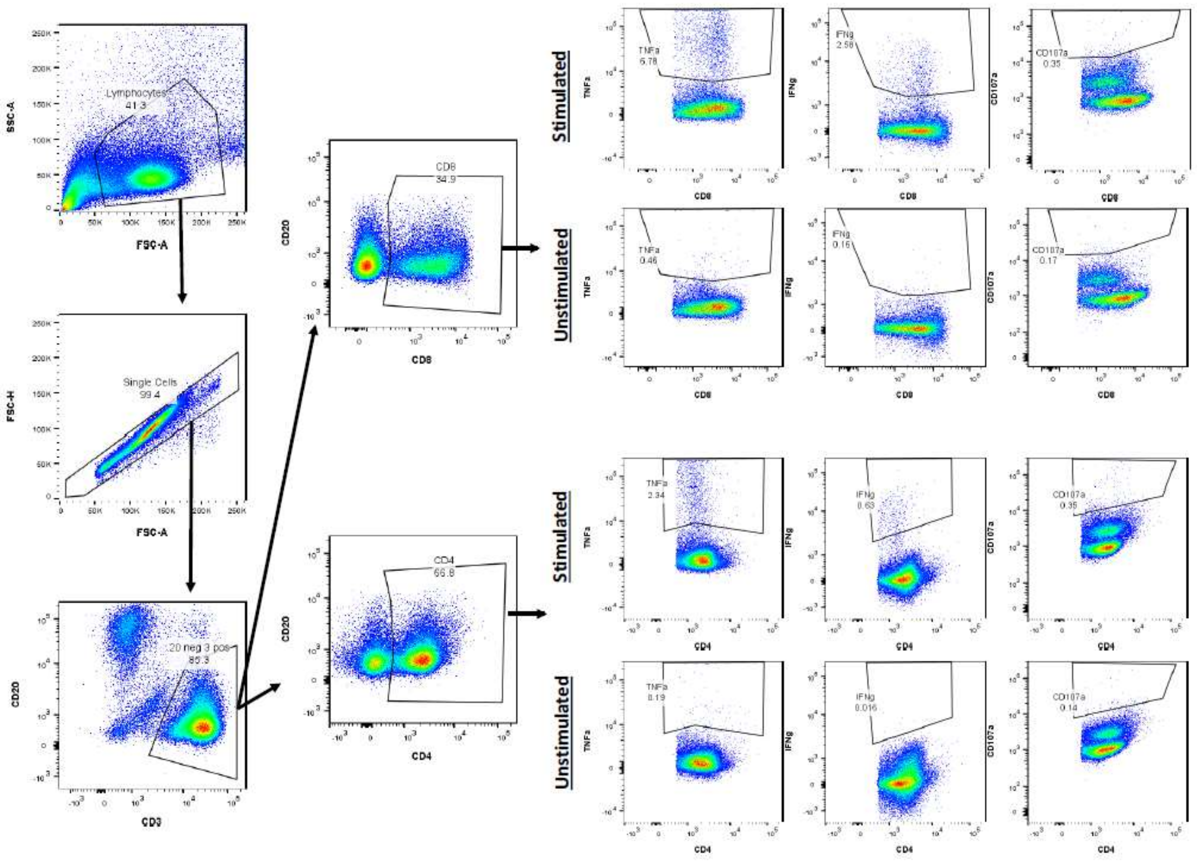


**S10 Figure. Gating strategy for B and T cells, and T cell response to stimuli.** Immunophenotyping of lymphocyte populations is shown. The resulting lymphocyte population was carried forward for selection of CD20+ CD3+ cells as B cells and CD3+ CD4+/ CD3+ CD8+ as CD4 T cells and CD8 T cells, respectively. Further gating is shown to represent immune functional responses measured after incubation of cells with DENV and ZIKV peptides.
